# Characterization of tRNA ligase function in pathogenic fungi and trypanosomes reveals the ligase domain as a promising drug target

**DOI:** 10.64898/2026.08.16.745097

**Authors:** Khondakar Sayef Ahammed, Pedro Miramon, Lukas Schrettenbrunner, Melissa R Cruz, Eun Young Huh, Huiqing Hu, Bhawana Israni, Hannah B. Wilson, Ziyin Li, Soo Chan Lee, Matthew G. Blango, Danielle A. Garsin, Michael C. Lorenz, Ambro van Hoof

## Abstract

The majority of eukaryotes encode some intron-containing pre-tRNAs. Splicing of these pre-tRNAs requires a dedicated tRNA splicing machinery. The fungal and trypanosome tRNA ligase, Trl1, and the human RNA ligase, RTCB, catalyze an essential step in tRNA splicing. However, Trl1 and RTCB are nonhomologous and biochemically and structurally distinct from each other. Therefore, Trl1 could serve as a broad-spectrum antifungal and anti-trypanosomal target. While the functions and requirements of the three catalytic Trl1 domains have been extensively characterized in the model yeast *Saccharomyces cerevisiae*, the roles of Trl1 orthologs in pathogenic fungi remain unexplored. Here, we validate Trl1 as one of the few promising novel drug targets for the development of antifungal therapeutics. Functional analyses of the three Trl1 domains show that only the “sealing” domain is essential for growth and viability in *Candida albicans* and *Aspergillus fumigatus*. In contrast, the two “healing” domains are dispensable in these pathogenic fungi, suggesting the presence of redundant healing enzymes, unlike in *S. cerevisiae*. These findings indicate that only the sealing domain is a good drug target. Our analysis also shows that the Mucor enzyme, which only contains the sealing domain, is essential. Using a *Caenorhabditis elegans* infection model of *C. albicans*, we further demonstrated that inhibiting Trl1 expression protects worms during an established infection. In contrast to these fungal pathogens, we show that all three domains of Trl1 are essential in *Trypanosoma brucei.* Our findings show that the essentiality of the Trl1 sealing is conserved in important human pathogens and provides an impetus for future drug development.

**SIGNIFICANCE:** Fungal infections are an important cause of human disease and death and difficult to treat and there is an urgent need to develop additional drugs. Based on studies in yeast, one promising target for antifungal drug development is the tRNA splicing pathway. Human tRNA ligase is fundamentally distinct from the fungal one. To investigate the possibility of developing tRNA ligase-targeting drugs, we investigated the function of the catalytic domains of fungal tRNA ligase in different fungal pathogens. Surprisingly, only the first domain is essential in these pathogens and yeast is not a good model fungus. In contrast, all three domains of Trypanosome tRNA ligase are essential. These findings provide an impetus for future drug development.

## INTRODUCTION

Fungal diseases are a threat, especially to the growing population of immunocompromised individuals. Globally, 1.5 million to 3.8 million deaths related to fungal diseases are reported annually (1, 2). In the United States, the Centers for Disease Control and Prevention (CDC) estimates that fungal diseases are associated with approximately 7,300 deaths, 130,000 hospitalizations, and 13 million outpatient visits each year, imposing an economic burden of $19 billion (3). Approximately 4.5% of all hospitalized patients in the US were prescribed a systemic antifungal during their stay between 2018 and 2023, an increase of 67% compared to 11 years earlier (4, 5).

The World Health Organization includes *Candida albicans, Candidozyma auris* (formerly *Candida auris*), *Cryptococcus neoformans*, and *Aspergillus fumigatus* in a “critical priority” group of the Fungal Priority Pathogens List (FPPL(6)). Collectively, these four pathogens are responsible for nearly half of fungal disease-related mortality in the United States and up to approximately 86% worldwide. In addition, Mucorales species remain significant causes of morbidity and mortality, and have very limited treatment options.

On the other hand, neglected tropical diseases caused by kinetoplastid parasites remain a major public health concern in tropical and subtropical regions of the world (7, 8). These include Human African Trypanosomiasis, Chagas disease, and leishmaniasis. Chagas disease, caused by *Trypanosoma cruzi*, is prevalent in Latin America and responsible for approximately 10,000 deaths annually, with an estimated 8 million people currently infected (9, 10). While cases of Human African Trypanosomiasis caused by *Trypanosoma brucei* have decreased in recent years, with nearly 1,000 new cases reported annually since 2018, the disease continues to pose a serious threat in several African countries (11).

Despite growing concerns about fungal and parasitic diseases, clinically approved treatment options remain limited. Currently, only three major classes of antifungal agents (polyenes, azoles, and echinocandins) are commonly used, and all three target the fungal cell envelope (12, 13). As of April 2025, the WHO reported nine antifungal agents in clinical trials (14), seven of which also target the fungal cell wall. Other drugs, such as Flucytosine (5-FC), have much more limited uses due to toxicity and the rapid emergence of resistance (15). Mucormycosis, caused by fungi of the order Mucorales, often requires surgical removal of the infected tissue or organ (16, 17), and azole resistance in *A. fumigatus* is a growing concern due to widespread use of azoles in agriculture (18-20). Expanding the range of molecular targets beyond the cell envelope is therefore critical to address increasing antifungal resistance. A few novel drug candidates for kinetoplastid diseases have shown encouraging results in clinical trials. For example, the proteasome inhibitor LXE408, the nitroimidazole prodrug Fexinidazole, and acoziborole analogs that target Cleavage and Polyadenylation Specificity Factor 3 (CPSF3) (21-23).

To identify additional antifungal drug targets, we developed a list of genes that have homologs in all invasive fungal pathogens, have no close homolog in humans, and are essential in all fungi where this has been tested (mostly *Saccharomyces cerevisiae*, *C. albicans*, *Schizosaccharomyces pombe*, and *C. neoformans*) (24-29). Pathogens are found across the diverse fungal kingdom. Due to this phylogenetic diversity within the fungal kingdom, and the close relationship with the animal kingdom, we identified only five candidate antifungal genes that fit these three criteria: *GSC1*, *FOL1*, *ALR1*, *FBA1*, and *TRL1* (Fig 1A). Gsc1 (a.k.a. Fks1) and Fol1 are the targets of current antifungal drugs: Gsc1 is the target of echinocandins and Fol1 is the target of sulfamethoxazole, which is used to treat *P. jirovecii* pneumonia (PCP) (30). Fungal tRNA ligase (Trl1) has been suggested as a promising antifungal drug target (31, 32). Trl1 enzymes from pathogenic fungi have been studied biochemically and structurally (31, 33-39), but little is known about its function other than being identified as an essential gene in high-throughput efforts (28, 29). Similarly, depleting *T. brucei* Trl1 prevents growth indicating that the enzyme is essential (40).

**Figure 1:**
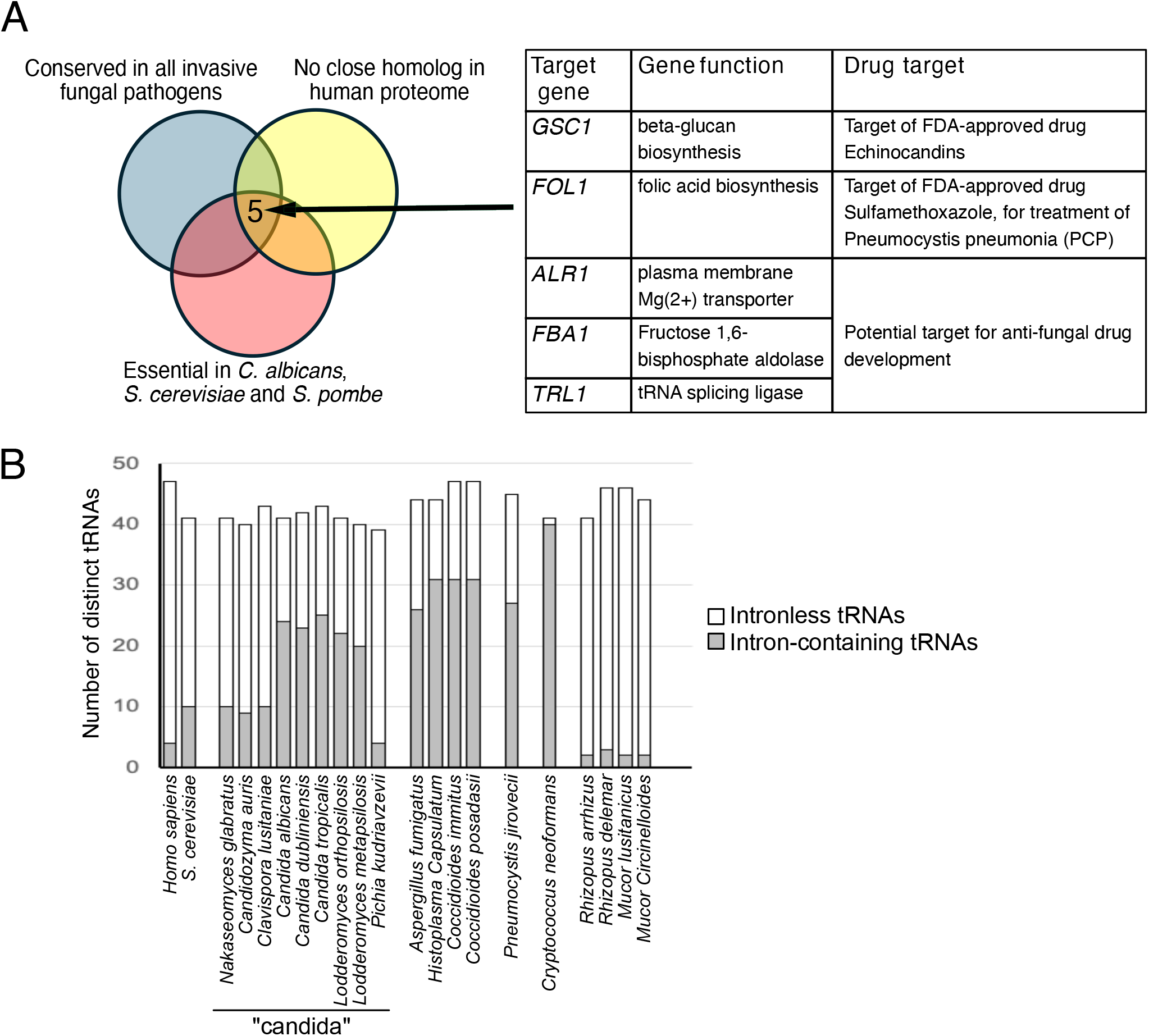
Bioinformatic analyses indicate that Trl1 is a promising drug target. (A) *Trl1* is one of five enzymes that are conserved in all invasive human pathogenic fungi examined, absent from human, and essential in all fungi where this has been tested. (B) Invasive human fungal pathogens have introns in the Tyr tRNA gene, the Ile tRNA gene with an UAU anticodon and a variable number of other tRNAs. “candida” indicates species that previously were included in the genus *Candida* but are currently in different genera and families than Candida

Trl1 and tRNA splicing have been extensively characterized in the model fungus, *S. cerevisiae*. The first step of the pre-tRNA splicing pathway is conserved among eukaryotes: cleavage of intron-containing pre-tRNAs by the tRNA splicing endonuclease (TSEN), resulting in a 5’ exon fragment with a 2’,3’ cyclic phosphate and a 3’ exon fragment with a 5’ OH end (Supplementary Figure 1). However, the subsequent step diverges between the “heal and seal pathway of *S. cerevisiae* and the “direct ligation” pathway of humans (and presumably fungi and animals). In humans, a single domain RNA ligase, RTCB, directly joins the 5’OH and 2’,3’ cyclic phosphate-containing exons to produce mature tRNAs (41-45). RTCB uses GTP to form an intermediate with GMP covalently bound to its active site histidine. Subsequently, this GMP is transferred to the 5’ exon to activate it (Supplementary Figure 1). In contrast, fungi employ a “heal and seal” pathway that uses a three-domain enzyme, Trl1, to execute consecutive “healing” and “sealing” reactions (34, 38, 46). In one healing reaction, the kinase domain of Trl1 uses GTP to phosphorylate the 5’ OH on the 3’ exon. In the second healing reaction, the cyclic phosphodiesterase (CPD) domain opens the 2’,3’ cyclic phosphate on the 5′ exon into a 2′-phosphate. Next, the N-terminal domain uses ATP to form an intermediate with AMP covalently bound to its active site lysine. Subsequently, this AMP is transferred to the 3’ exon to activate it. Both the N-terminal domain and the full-length protein are named ligase. We refer to the N-terminal domain as the “sealing domain” to avoid confusion. Thus, the nucleotide used (GTP vs ATP), the catalytic residue (Lys vs. His), the exon that is activated (3’ vs 5’), the product (with 2’ phosphate, versus without it), and the protein structure are all fundamentally different between Trl1 and RTCB. Due to these fundamental differences, Trl1 was proposed as an attractive target for the development of antifungal drugs (31, 47). However, an understanding of Trl1 functions in pathogenic fungi species is lacking.

Although the name-sake function of tRNA ligases (Trl1 and RTCB) is to ligate tRNA exons, the enzymes have an important second function during unfolded protein stress. In the Unfolded Protein Response pathway, the endoribonuclease Ire1 cleaves a specific mRNA (Hac1 in fungi, Xbp1 in animals) in two places, releasing an atypical intron. The 5’ and 3’ fragments are then ligated by Trl1 or RTCB to create an mRNA that encodes a transcription factor that restores proteostasis. The biochemistry and mechanism of this *HAC1/XBP1* splicing is identical to the tRNA splicing mechanism described above. This UPR is generally not essential, but Ire1 and Hac1 mutants in several fungi have defects in pathogenesis (48-50). Although Trl1 has functions in both tRNA splicing and UPR, only the tRNA splicing function is essential in *S. cerevisiae* and *T. brucei* (40, 51), as shown by the viability of Trl1 deletion/depletion if all tRNAs are artificially expressed from intronless genes. Nevertheless, the importance of the UPR for fungal pathogenesis further strengthens the promise of Trl1 as an antifungal drug.

Previous work from us and others identified conserved Trl1 orthologs across major human pathogenic fungal species (31, 33-39). In this study, we experimentally validate the sealing domain of Trl1 as a strong antifungal drug target. We show that this domain of Trl1 meets key criteria for an effective antifungal target, as it is essential for growth in the fungal pathogens *C. albicans*, *A. fumigatus*, and Mucor, and its inhibition is fungicidal in S. cerevisiae. Genetic inhibition of Trl1 in *C. albicans* can mitigate an established animal infection. Remarkably, the healing domains of Trl1 are not essential in pathogenic fungi, suggesting that fungi typically contain redundant healing activities that have been lost in *S. cerevisiae*. In contrast, our results show that in the parasite *Trypanosoma brucei* all three catalytic domains are essential, indicating that each domain may represent a viable drug target in trypanosomes. Together, these findings highlight functional variations of the tRNA splicing pathway across fungi and trypanosomes and validate the Trl1 sealing domain as a promising target for the development of dual-use antifungal and antiparasitic drugs.

## RESULTS

### Inhibition of *Trl1* in the model fungus *S. cerevisiae* is fungicidal

Our analysis of conserved essential fungal enzymes without human orthologs identified two current drug targets (Gsc1 and Fol1) and three potential novel targets (Figure 1A). Among these, Alr1 and Fba1 have been explored somewhat in fungal pathogens (52, 53), while the function of Trl1 in pathogens is largely unexplored. We therefore focused on Trl1. We first surveyed the tRNA intron content of major fungal pathogens, and showed that they all contained an intron in the Tyr tRNA and one of the two Ile tRNAs. Some fungal pathogens (e.g. *Mucor*) contained no other tRNA introns, while others contained an intron in most tRNA genes (Figure 1B). For example, *C. neoformans* requires splicing to translate all codons for 16 or 17 amino acids (for *C. neoformans var. grubii H99* and *C. neoformans var. neoformans JEC21*, respectively) and some codons for the others. Without tRNA splicing *C. neoformans* can only translate six or seven codons (CAU, CAC, GGA, GGC, GGG and CAG for H99 and also AUG for JEC21). In addition, neither strain can initiate translation without tRNA splicing because the initiator tRNA genes each contain an intron. All of these fungal pathogen introns resemble the tRNA introns of *S. cerevisiae* in length and in being one base 3’ of the anticodon. Other pathogenic fungi have a range of tRNA introns between the *Mucor* and *Cryptococcus* extremes.

An ideal antifungal drug is fungicidal, meaning it kills fungi instead of merely inhibiting their growth. A major drawback of the azole drugs is that they are largely fungistatic. We therefore used tools available in *S. cerevisiae*, but not in fungal pathogens, to ask whether targeting Trl1 is fungicidal or fungistatic. Although we could not find a universal definition of “fungicidal”, a common criterium is that the drug causes a 10-fold reduction in viable cells within 24 hours. We used a temperature-sensitive strain of *S. cerevisiae* (*trl1-ts*) that grows normally at room temperature but is defective in growth at higher temperatures (24, 54). The *trl1-ts* strain was grown at room temperature and then incubated at the restrictive temperature (37^°^C) for 0 to 24 hours to inhibit Trl1. Following this incubation, the cultures were plated and incubated at room temperature for 5-6 days to measure the number of colony-forming units (Figure 2A). As a control, we incubated wild-type *S. cerevisiae* with 5 µg/mL nystatin, which is known to be fungicidal. We observed a 10-fold decrease in CFU after 16 hours of nystatin treatment, consistent with its known fungicidal activity (Figure 2B-C) (55). In comparison, inactivation of Trl1 resulted in a 10-fold decrease in CFU as soon as 8 hours. This suggests that repressing Trl1 activity not only restricts the growth of *S. cerevisiae* but also reduces cell viability. The few *trl1-ts* colonies that arose after inactivation of Trl1 for 24 hours remained temperature-sensitive, indicating they are not due to suppressors/resistance mechanisms. We conclude that a Trl1 inhibitor would likely be fungicidal, as Trl1 inhibition outperformed nystatin treatment.

**Figure 2:**
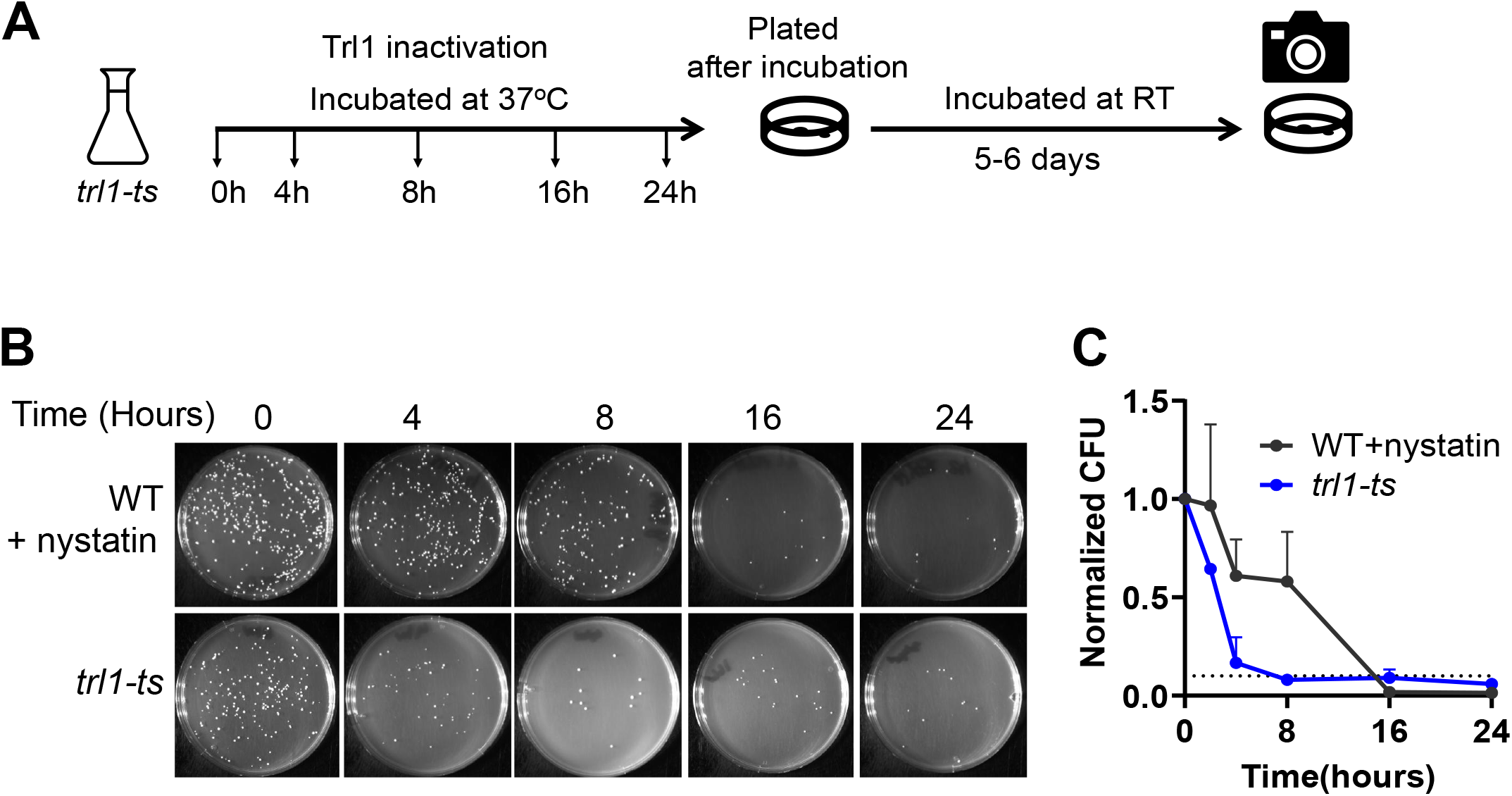
Inhibition of Trl1 in the model fungus *S. cerevisiae* is fungicidal. (A) Schematic of the experimental design. The *trl1-ts* strain was grown at room temperature to mid-log phase and incubated at 37°C at the indicated time periods and plated on YPD for 5 days. After 5 days images were captured and colonies were counted. (B) The *trl1-ts* strain resulted in a drastic reduction in colony-forming units (CFU) when incubated at the restrictive temperature, indicating that Trl1 inhibition is fungicidal. As a control, wild-type *S. cerevisiae* was treated with 5 µg/mL of the fungicidal drug nystatin. (C) The number of CFU remaining at the indicated timepoints was normalized to the colony count at 0 hours of incubation. The average CFU and standard deviation from two replicates is plotted.

### Trl1 is essential in the fungal pathogen *Candida albicans*

An important requirement for an antifungal drug target is that it be essential for viability in major fungal pathogens. High-throughput genome-wide studies have concluded that the Trl1 orthologs of *C. albicans* and *C. neoformans* are essential (28, 29). The *C. albicans* study used random transposon insertions into a stable haploid derivative strain. Insertions into *TRL1* were underrepresented, suggesting that *TRL1* is an essential gene. We sought to confirm the essentiality in the commonly used diploid strain SC5314 because a few transposon inserts were obtained, and haploid strains are not found in a natural or clinical setting. Although we and others routinely knock out nonessential genes in SC5314, we failed to obtain homozygous *trl1Δ/trl1Δ* deletion strains and instead recovered >100 heterozygous *TRL1/trl1Δ* transformants. However, when a complementing *TRL1* gene was first integrated in a heterologous locus of the genome, we obtained 11 knockouts at the endogenous locus and 13 heterozygote *TRL1/trl1Δ*. The ability to obtain *trl1Δ/trl1Δ* homozygotes only after integration of *TRL1* at a complementing locus suggests that *TRL1* is indeed essential.

To validate the essentiality of *TRL1*, we used a doxycycline-repressible TET-off promoter. Specifically, we created a heterozygous conditional mutant of *C. albicans* by deleting one *TRL1* allele with a *FLP1-SAT1* cassette and replacing the promoter of the other allele with a TET-off promoter using CRISPR-CAS9-targeted integration (Figure 3A) (56-58). This conditional *C. albicans* mutant *(PtetO-TRL1/trl1Δ*) and the intermediate heterozygote (*TRL1/trl1Δ*) grew at close to wild-type rates in media without doxycycline. Upon exposure to doxycycline, the *C. albicans* mutant (*PtetO-TRL1/trl1Δ*) showed a strong growth defect, confirming that *TRL1* is required for growth. To further validate this conclusion, we reintroduced the wild-type *C. albicans TRL1* gene into the conditional mutant at the *NEUT5L* neutral locus by homologous recombination (*PtetO-TRL1/trl1Δ+TRL1)*. This expression of Trl1 from a different locus fully rescued the growth defect caused by Trl1 repression. Therefore, Trl1 is essential in *C. albicans* and a valid drug target.

**Figure 3:**
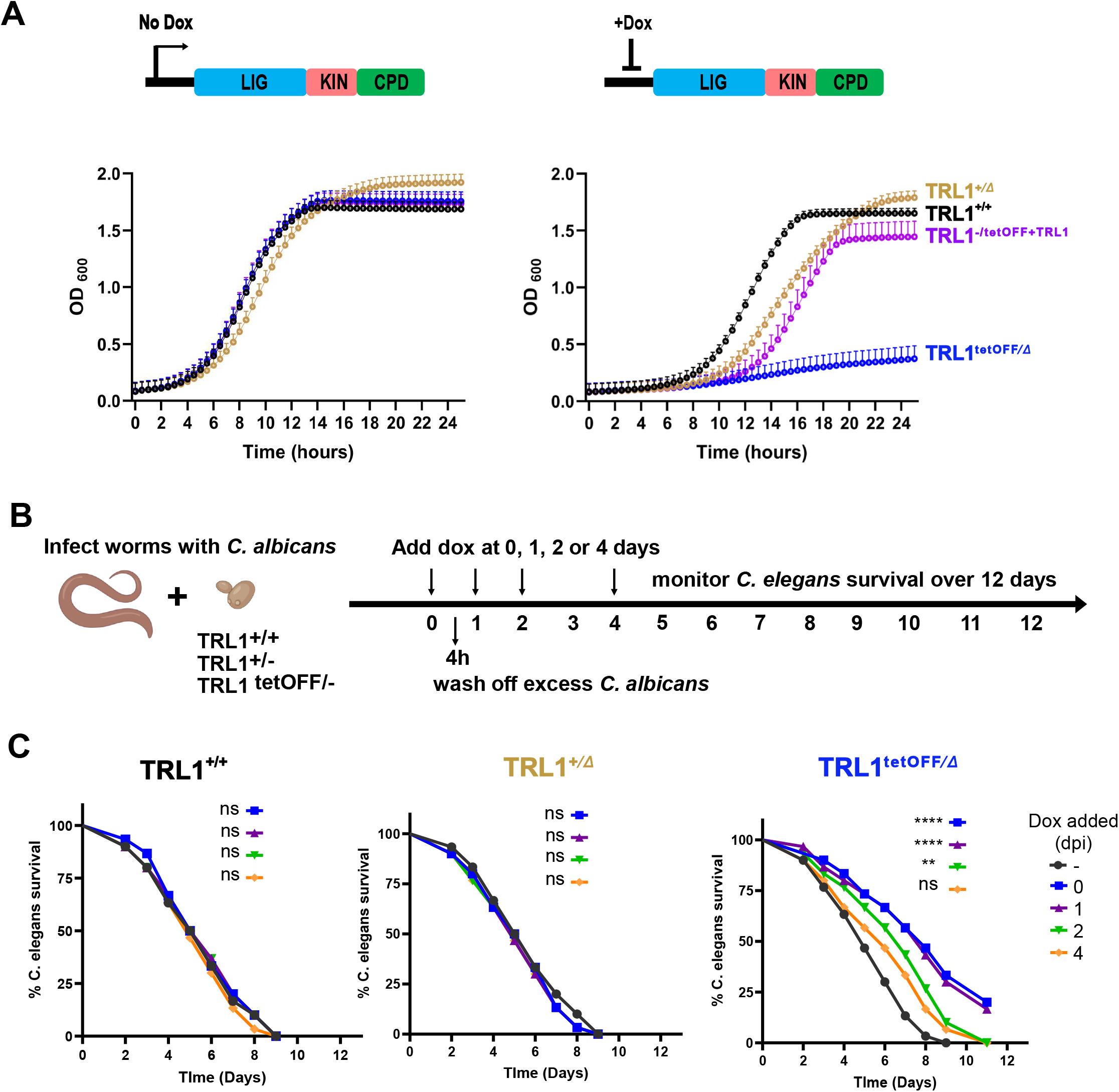
*C. albicans* Trl1 is essential. (A) A doxycycline repressible conditional *TRL1* mutant confirms that *TRL1* is essential in *C. albicans*. In the presence of doxycycline (right panel), the conditional mutant (*PtetO-TRL1/trl1Δ*; blue graph) showed impaired growth compared to the wild type (TRL1^+/+^; black) or heterozygous mutant (*TRL1/TRL1Δ*; brown). Growth is restored by an ectopic copy of *TRL1* at the *NEUT5L* locus (*PtetO-TRL1/trl1Δ+TRL1*: purple graph). In the absence of doxycycline all three strains grew similarly to each other (left panel). (B) Schematic representation of the *C. elegans* killing assay with the *C. albicans* conditional mutant of *TRL1* (*PtetO-TRL1/trl1Δ*). The indicated *C. albicans* strains were incubated with wild type N2 *C. elegans* for 4 hours to establish an infection. Excess *C. albicans* was removed and doxycycline was added at the indicated time points in order to repress Trl1 expression in the *PtetO-TRL1/trl1Δ* strain. Survival of animals was monitored over an 11-day period. (C) The percentage of viable animals was plotted for each strain of *C. albicans*. Infection by the wild type (*TRL1^+/+^*; left panel) and heterozygous mutant (*TRL1/TRL1Δ*; middle panel) was unaffected by doxycycline. The survival of animals after infection by the conditional mutant (*TRL^/Δ/TEToff^*; right panel) was improved after doxycycline treatment with early treatment being more effective than later treatment when animals were already dying from the infection. Shown is one of three biological replicate experiments. Statistical significance using log-rank analysis was assessed by comparing the doxycycline-treated survival curves to the untreated control. p values are indicated ns= not significant

### Inhibition of *TRL1* can treat an established *C. albicans* infection in an animal model

Although *TRL1* meets the major criteria for an antifungal drug target, it remains unclear whether inhibition of *TRL1* improves host survival during fungal infection in an animal model. We hypothesized that repression of *TRL1* in the doxycycline-repressible conditional mutant *(PtetO-TRL1/trl1Δ)* would confer a survival advantage in a *Caenorhabditis elegans* model infected with *C. albicans*. To test this assumption, we used a killing assay in which animals were infected with wild-type *C. albicans*, a heterozygous mutant (*TRL1/trl1Δ*), or a doxycycline-repressible conditional *trl1* mutant (*PtetO-TRL1/trl1Δ*) during a 4-hour exposure. Excess *C. albicans* cells were then washed off. Doxycycline was added either at the start of the infection or 1, 2, or 4 days later. (Figure 3B). The survival of the animals was monitored over the following 12 days. Importantly, repression of Trl1 expression in the conditional mutant (*PtetO-TRL1/trl1Δ*) upon doxycycline exposure protected animals from *C. albicans* infection (Figure 3C). The increase in animal survival correlated with the timing of doxycycline treatment. When *TRL1* was repressed on the day of infection or one day post-infection, animal survival was significantly improved compared with animals without *TRL1* repression in all biological replicates. Although animals in which *TRL1* was repressed on day 2 or day 4 also showed a trend to a survival advantage relative to the no-doxycycline control, the effect was less pronounced and not statistically significant in all replicates. We anticipate that a Trl1-inhibiting drug would inhibit Trl1 activity more rapidly than depleting the mRNA and protein after transcription shut-off, therefore improving efficacy. In controls, animals infected with wild-type or heterozygous *TRL1/trl1Δ* mutant *C. albicans* exhibited similar killing kinetics after doxycycline exposure, indicating that doxycycline acted through the intended mechanism and not through off-target mechanisms. As an additional control, all three strains were equally pathogenic in the absence of doxycycline. These results suggest that inhibition of Trl1 with inhibitors could treat an already established infection.

### Only the sealing activity of *C. albicans* Trl1 is essential

Fungal Trl1 is a three-domain enzyme that carries out three consecutive reactions of the tRNA exon ligation pathway (supplemental figure 1). In principle, each of the domains would make a good antifungal target, as each of the activities is required for tRNA splicing and all three domains are essential in *S. cerevisiae*. To test whether the catalytic activity of all three domains of *C. albicans* Trl1 is essential for survival, we identified the key catalytic amino acid residues within the three domains of *C. albicans*. (59) (Figure 4A). In the sealing domain, the catalytic lysine that forms a covalent intermediate with AMP is lysine 108. Within the P-loop of the kinase domain substitution of critical Lys and Thr residues abolishes catalytic activity in *S. cerevisiae* (59). The corresponding residues in *C. albicans* Trl1 are K425 and T426. In the CPD domain a His residue is indispensable for catalytic activity and the corresponding residue in *C. albicans* Trl1 is H767 (59). To determine whether the designed K108A, K425A T426A, and H767A mutations in *C. albicans* Trl1 indeed inactivate the corresponding domains, we expressed the mutant *C. albicans* proteins in *S. cerevisiae.* Each of the three mutant *C. albicans* Trl1 proteins failed to complement the lethality caused by deletion of *S. cerevisiae TRL1* (Figure 4B). These results confirm that the mutations abolish the function of the respective domains of *C. albicans* Trl1.

**Figure 4:**
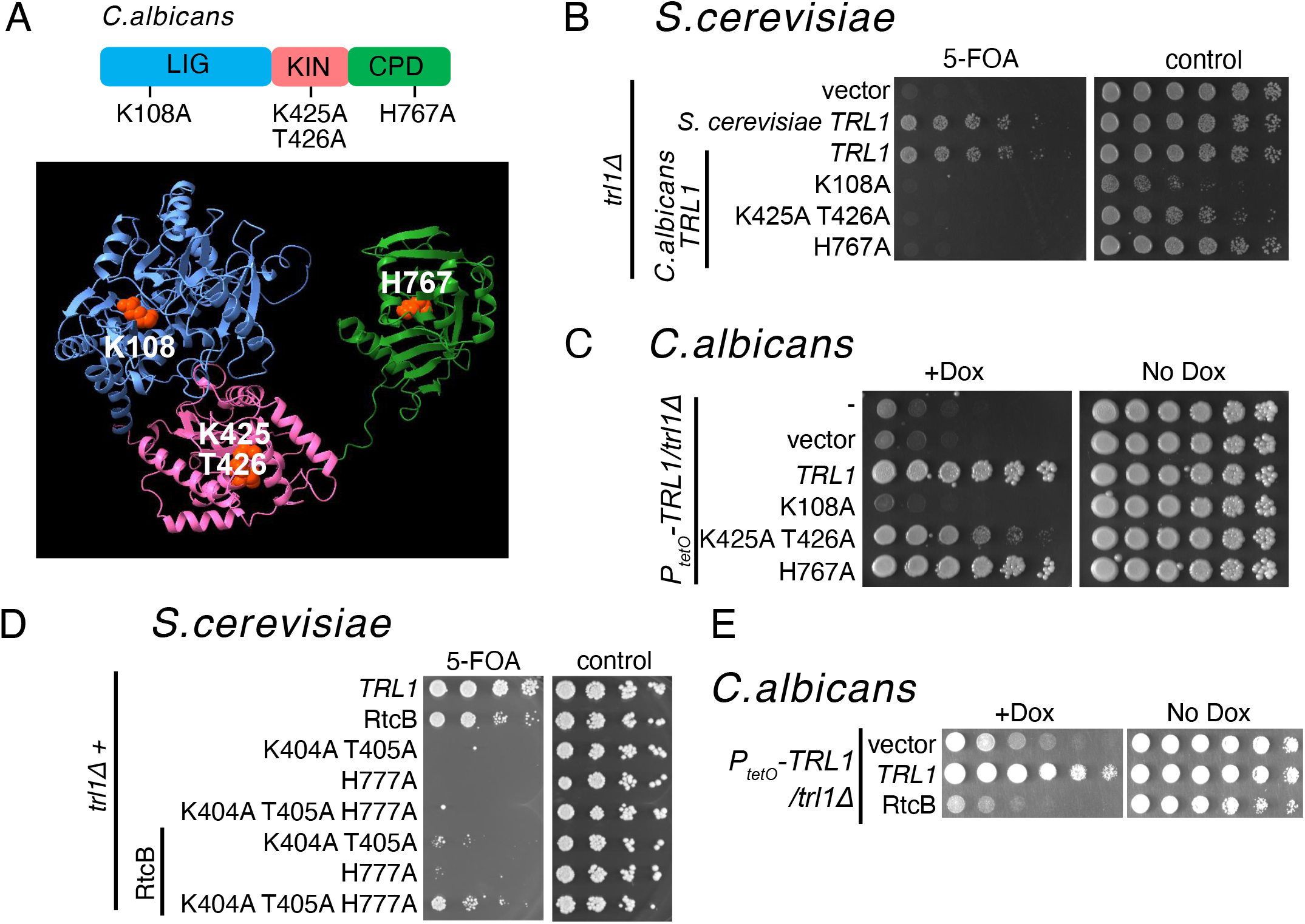
Only the sealing activity of *C. albicans* Trl1 is essential. (A) The active-site mutations in ligase (K108A), kinase (K425A T426A), and CPD (H767A) domains of *C. albicans* Trl1 corresponding to the *S. cerevisiae* Trl1 mutations that inactivate the function of individual domains. Shown is an AlphaFold-generated structure of *C. albicans* Trl1 depicting three individual domains: ligase (blue), kinase (pink), and CPD (green). The active-site residues, represented as red spheres, are mutated to alanine for functional analysis. (B) All three activities of *C. albicans* Trl1 are required to complement *S. cerevisiae trl1Δ*. Shown is a plasmid shuffle assay in a *S. cerevisiae trl1Δ* strain complemented with a *TRL1, URA3* plasmid. Serially diluted *S. cerevisiae* also cells expressing either the wild-type *C. albicans* Trl1 or the indicated mutants, are spotted on SC media lacking leucine and histidine (SC-Leu-His) as a control. The same cells were spotted on media containing 5-FOA, where growth indicates that the URA plasmid can be lost because the *C. albicans* Trl1 complements. (C) Only the ligase activity of Trl1 is essential in *C. albicans.* The three catalytic mutants were ectopically expressed in the TET-repressible conditional mutant. Serially diluted cells were spotted on YPD media with or without doxycycline. (D) *E. coli* RtcB can complement S. cerevisiae *trl1Δ* (row 2), but poorly complements Trl1 with a mutation in one of the healing domains (rows 6 and 7). However, the combination of mutations in both healing domains can be rescued by expressing RtcB (row 8). (E) *E. coli* RtcB fails to complement conditionally repressed *TRL1* in *C. albicans*. The *E. coli* RtcB was expressed in the doxycycline-repressible *TRL1 C. albicans* strain background. The growth defect caused by *TRL1* repression can be rescued by expressing a wild-type *C. albicans* Trl1, but not RtcB, suggesting that alternative healers in *C. albicans* modify tRNA ends that cannot be ligated by RtcB.

We next expressed the *C. albicans* Trl1 mutants from the *NEUT5L* locus in the doxycycline-repressible conditional *trl1* mutant (*PtetO-TRL1/trl1Δ*) to determine which Trl1 domains are essential in *C. albicans*. The strain expressing the sealing domain mutant (*PtetO-TRL1/trl1Δ+trl1-K108A*) failed to grow upon exposure to doxycycline. Thus, the sealing activity of *C. albicans* Trl1 is essential as expected (Figure 4C). Surprisingly, the Trl1-kinase mutant (*PtetO-TRL1/trl1Δ+trl1-K425AT426A*) and CPD mutant (*PtetO-TRL1/trl1Δ+trl1-H767A*) grew when endogenous *TRL1* was repressed by doxycycline. Notably, the Trl1-kinase mutant grew slowly instead of fully complementing the growth defect caused by Trl1 repression. However, the CPD mutant complementation restored growth to near wild-type levels. Because we did not expect complementation by the healing domain mutants, we ensured the strains were correct by long-read whole-genome sequencing. Thus, the healing activities of *C. albicans* Trl1 are dispensable. The *Saccharomyces trl1Δ* complementation results (Figure 4B) indicate that splicing by *C. albicans* Trl1 requires healing activities. We conclude that *C. albicans* probably expresses alternative healing activities that are redundant with those of Trl1. An attempt to identify potentially redundant healing enzymes in *C. albicans* did not reveal any close paralogs of the kinase or CPD domain, suggesting that the redundant healing activities in *C. albicans* are not closely related to the healing domains of *S. cerevisiae* Trl1 (see discussion). Consistent with our conclusion, it has previously been shown that non-orthologous healing activities from bacteriophage T4 or mammals can complement Trl1 defects in *S. cerevisiae* (60-62). Therefore, the ligase domain of Trl1 appears to be the most suitable target for designing broad-spectrum antifungal drugs.

To further test the presence of redundant healing activities, we expressed the *E. coli* RNA ligase RtcB in *S. cerevisiae* and *C. albicans*. RtcB expression can rescue the *trl1Δ* growth defect phenotype in *S. cerevisiae* (63) by directly ligating the 2’,3’ cyclic phosphate and 5’ OH ends created by TSEN. We predicted that in the presence of functional healing activities, the 2’,3’ cyclic phosphate and 5’ OH ends would no longer be available for RtcB. To test this, we first expressed RtcB in *S. cerevisiae* with point mutations in one of the healing activities. The strain with the kinase point mutation should accumulate 2’phosphate ends instead of 2’,3’ cyclic phosphate ends, and as expected, RtcB complements this strain poorly compared to a *trl1Δ* strain. Similarly, the strain with the CPD point mutation should accumulate 5’ phosphate ends instead of 5’ OH ends, and as expected, RtcB complements this strain poorly. Strikingly, a *S. cerevisiae* strain with defects in both healing domains could be complemented by expressing RtcB, consistent with the accumulation of 5’ OH and 2’,3’ cyclic phosphate ends. (Figure 4D). Collectively, these results confirm that RtcB complementation is impaired by the healing activities. We next analyzed whether RtcB can complement Trl1 depletion in *C. albicans* (Figure 4E). Consistent with the presence of alternative healing activities, RtcB did not restore growth upon depletion of *C. albicans* Trl1. Although other possibilities for this failure to complement cannot be ruled out, this finding supports the hypothesis that *C. albicans* possesses alternative healing activities that are redundant with those of Trl1.

Based on our results that repression of Trl1 protects *C. elegans* during a *C. albicans* infection, we investigated whether loss-of-function mutations in different domains of *C. albicans* Trl1 have distinct effects on host survival during infection. We performed a *C. elegans* killing assay using a doxycycline-repressible conditional *trl1* mutant (*PtetO-TRL1/trl1Δ*) that expressed either wild-type or mutant Trl1 from the *NEUT5L* locus (Figure 4). As an additional negative control, the conditional *trl1* mutant was also transformed with an empty vector integrated at the *NEUT5L* locus. As expected from our in vitro growth analysis, Trl1 mutants with defects in one of the healing domains complemented *PtetO-TRL1/trl1Δ* during this animal infection, as did the wild-type enzyme. (Figure 5). In contrast, ectopic expression of the sealing domain mutant failed to complement (Figure 5). Together, our results suggest that the sealing domain (but not the healing domains) of *C. albicans* Trl1 is a promising antifungal drug target. Our data also explain why random transposon insertions in *C. albicans TRL1* were rare, with only two insertions reported: one in codon 406, in the linker between the sealing and kinase domains, and one in codon 630, in the linker between the kinase and CPD domains (28). Thus, both transposon insertions allow expression of the sealing domain.

**Figure 5:**
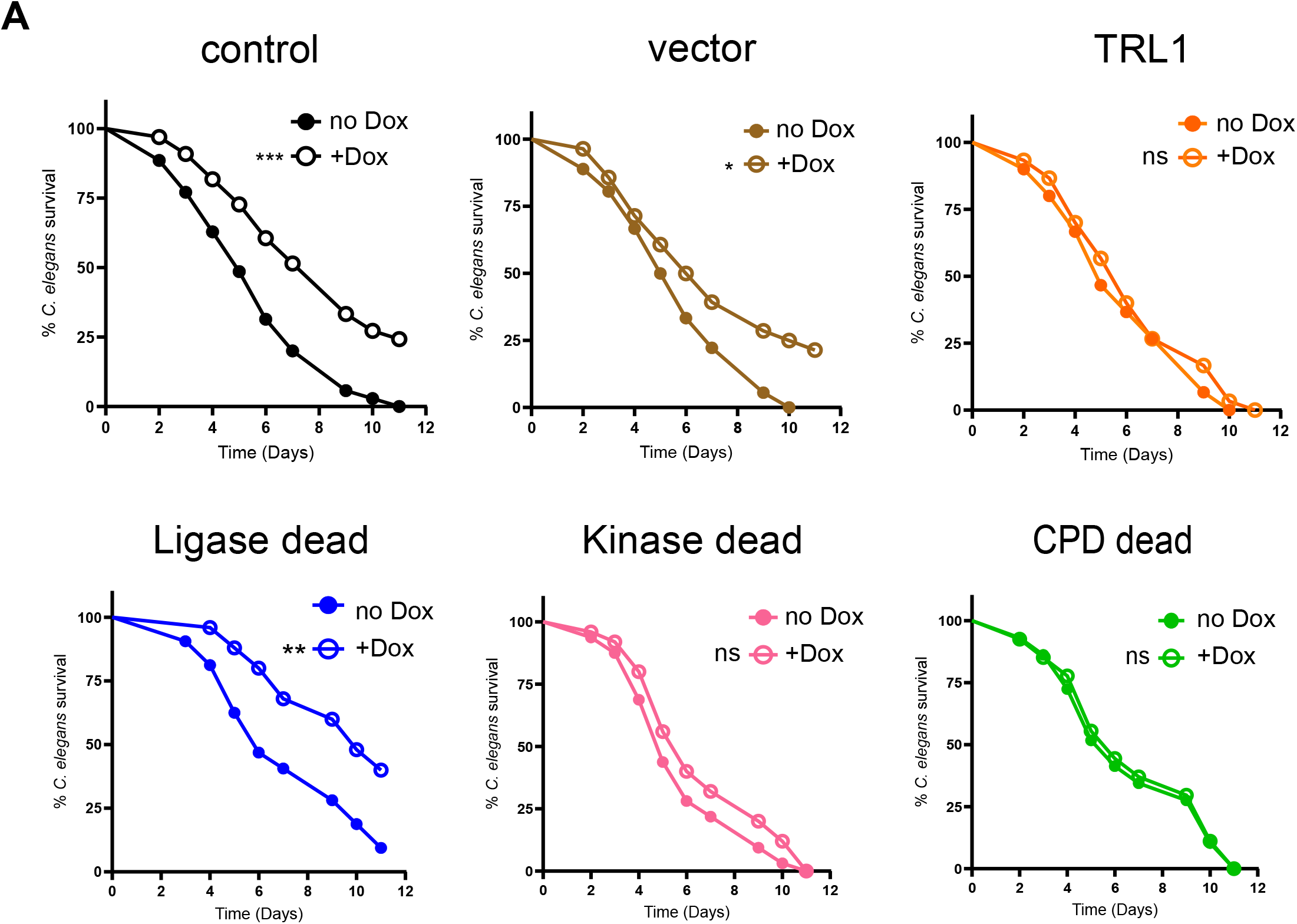
The ligase activity of *C. albicans* Trl1 is required during an animal infection. *C. elegans* killing assays were performed and analyzed with the indicated strains as described for figure 3C. +dox indicates that doxycycline (100 µg/mL) was added at the start of infection. Shown is one of three biological replicate experiments. Statistical significance using log-rank analysis was assessed by comparing the doxycycline-treated survival curves to the untreated control. p values are indicated ns= not significant

### Only the sealing activity of *Aspergillus fumigatus* Trl1 is essential

The surprising differences in domain requirements between *S. cerevisiae* and *C. albicans* prompted us to examine *A. fumigatus*. It is a filamentous fungus listed as one of the four critical priority fungal pathogens on the WHO Fungal Priority Pathogens List, and like *C. albicans* and *S. cerevisiae* is a member of the phylum Ascomycota. *Aspergillus* is in a different subphylum than *Candida* and *Saccharomyces* and diverged from these yeasts approximately 500 million years ago. We and others have previously shown that *A. fumigatus* encodes a canonical Trl1 ortholog with three distinct domains (Figure 6A) (31, 32). Currently, it is unclear whether *A. fumigatus* Trl1 is essential for viability, although the presence of tRNAs with introns suggests that tRNA splicing is essential (Figure 1B). To test this, we replaced the native *trl1* promoter with a doxycycline-inducible promoter (*TRL1^TETon^*) to conditionally express Trl1 (Figure 6). The conditional mutant showed a growth rate similar to that of wild-type *A. fumigatus* in media containing doxycycline (Figure 6). Growth was severely impaired after 48 hours on media without doxycycline. To further confirm that the growth defect of the conditional *trl1* mutant is solely because of the loss of *trl1*, we inserted *A. fumigatus trl1+* under the control of its native promoter, into a genomic ‘safe haven’ region (64). This extra *trl1+* complemented the *trl1^TETon^*strains, confirming that *A. fumigatus* Trl1 is essential for viability.

**Figure 6:**
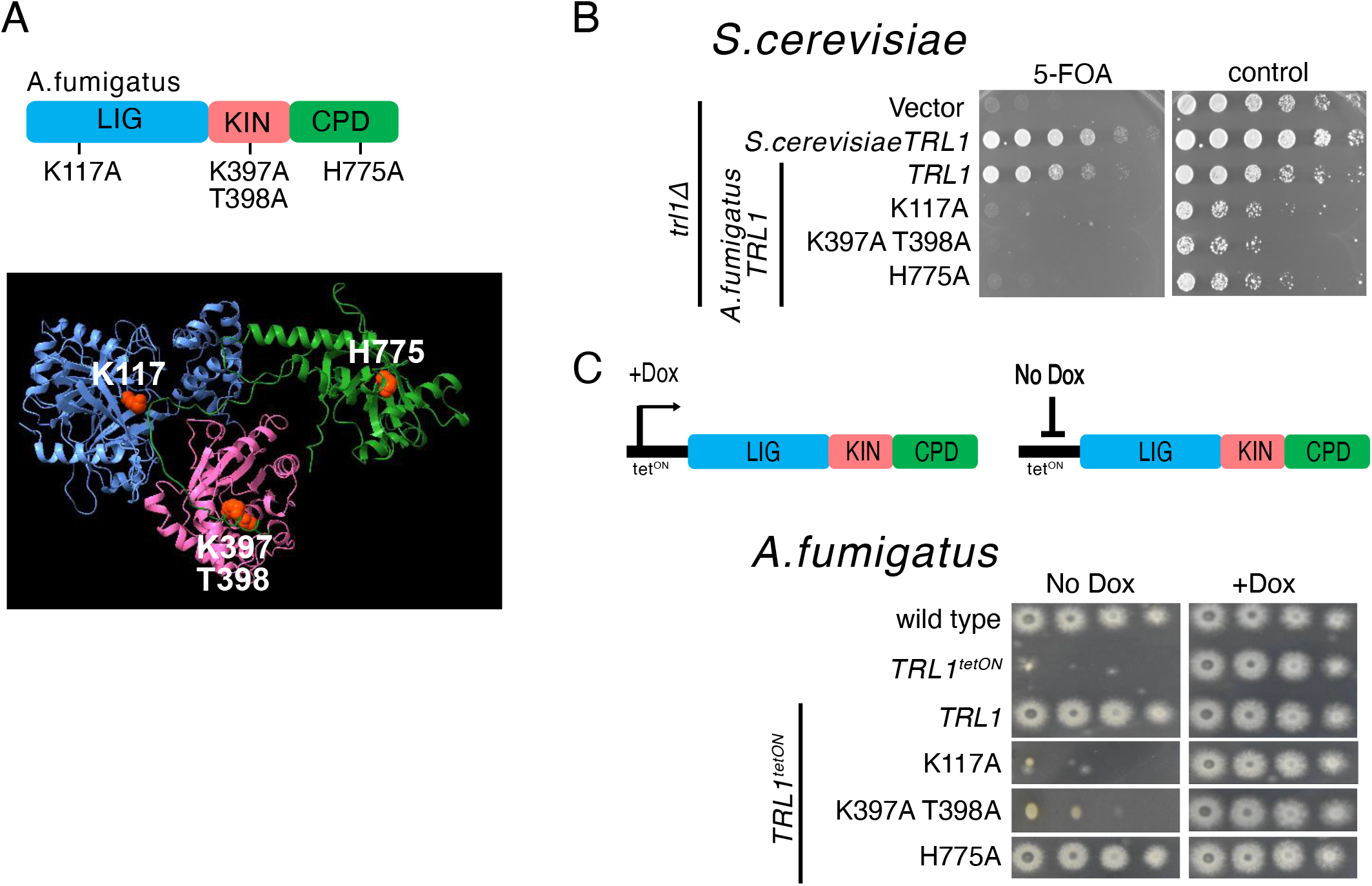
The sealing domain of Trl1 is essential in *Aspergillus fumigatus.* (A) Schematic and AlphaFold predicted structure of *A. fumigatus* Trl1 with the three active sites indicated as in figure 4A. (B) All three activities of *A. fumigatus* Trl1 are required to complement *S. cerevisiae trl1Δ*. The experiment was performed as in Figure 4B. (C) An *A. fumigatus* strain with a doxycycline inducible *trl1* gene grows in the presence of doxycycline, but not in its absence (row 2). This growth defect is complemented by ectopic expression of wild-type Trl1 (row 3) or the CPD domain mutant (row 6). The *A. fumigatus* Trl1 with a defective sealing domain fails to complement (row 4) and the kinase mutant complements very poorly (row 5). Strains were serially diluted and plated on AMM with or without 3 µg/mL doxycycline, and growth was recorded at 48 hours.

To test which activities of Trl1 are essential in *A. fumigatus* we mutated critical amino acid residues in the sealing (K117), kinase (K397, T398), and CPD (H775) domains based on sequence alignments and AlphaFold-predicted structures (Figure 6A). All three *A. fumigatus* Trl1 mutants failed to complement the growth defect of *S. cerevisiae trl1Δ*, indicating that these point mutations disrupt the activity of their respective domains as designed (Figure 6B). When expressed from the safe haven locus in the *trl1^Tet-ON^* conditional mutant strain, in the absence of doxycycline, the strain expressing the sealing domain mutant (K117A) exhibited severely reduced growth, indicating that the sealing domain is essential for the viability of *A. fumigatus* (Figure 6C). The strain expressing the kinase defective mutant showed slightly better growth, but also was severely impaired. Like we observed in *C. albicans*, the CPD domain mutant grew well.

These results parallel the Trl1 requirements observed in *C. albicans:* the sealing activity is essential in both species, the CPD domain is completely dispensable, and loss of the kinase activity results in slow growth in both species (although to a more dramatic extent in *Aspergillus*). Nevertheless, the strict requirement for the Trl1 ligase domain highlights this domain as a promising antifungal drug target. In contrast, targeting the healing domains may be less effective, as both domains appear dispensable for viability in the two fungal pathogens tested, presumably because they express redundant healing activities.

### The ligase-only Trl1 of *Mucor circinelloides* is essential

Human pathogenic fungal species from the early-diverging order Mucorales, including pathogens of the *Mucor* and *Rhizopus* genera, encode a truncated Trl1 ortholog that contains only the sealing domain. These fungi diverged from other fungal pathogens approximately 700 million years ago. Previous studies have shown that the truncated Mucorales Trl1 can complement the sealing function of *S. cerevisiae* Trl1 (32, 33). However, it remains unclear whether this “sealing-only” Trl1 is essential for the viability of Mucorales species. The genetic tools available for Mucorales are limited, including that no facile inducible or repressible promoter is available (65). Furthermore, Mucorales grow as coenocytic hyphae with many haploid nuclei in a single contiguous cytoplasm. Because of the multinucleated nature, essential genes can be disrupted in *Mucor*, but only in the presence of a wild-type nucleus in the same shared cytoplasm. Although *M. circinelloides* spores are also multi-nucleate, it also produces a subpopulation of smaller (3µm) spores with one or two nuclei. The inability to recover small uni-nucleate mutant spores is indicative of gene essentiality (65).

To test the essentiality of the *M. circinelloides trl1* gene, we disrupted it with the *pyrG-dpl237* marker (Figure 7A, B; *pyrG* is the ortholog of *S. cerevisiae URA3*). We obtained 4 different *trl1Δ::pyrG* multi-nucleate transformants that were confirmed by PCR amplification across the 5′ and 3′ junctions of the deletion cassette (Figure 7B). However, in each case a wild-type *trl1+* gene remained, suggesting these strains were heterokaryotic (hyphae with mix of mutant and wild-type nuclei). Mononucleated small Ura+ spores were enriched by filtration through a 5 µm filter and analyzed by PCR. Analysis of 21 small spore progeny from two independent transformants showed that each progenitor retained nuclei with an intact *TRL1* gene (Figure 7C). We conclude that the *M. circinelloides* sealing-only Trl1 is essential for viability.

**Figure 7:**
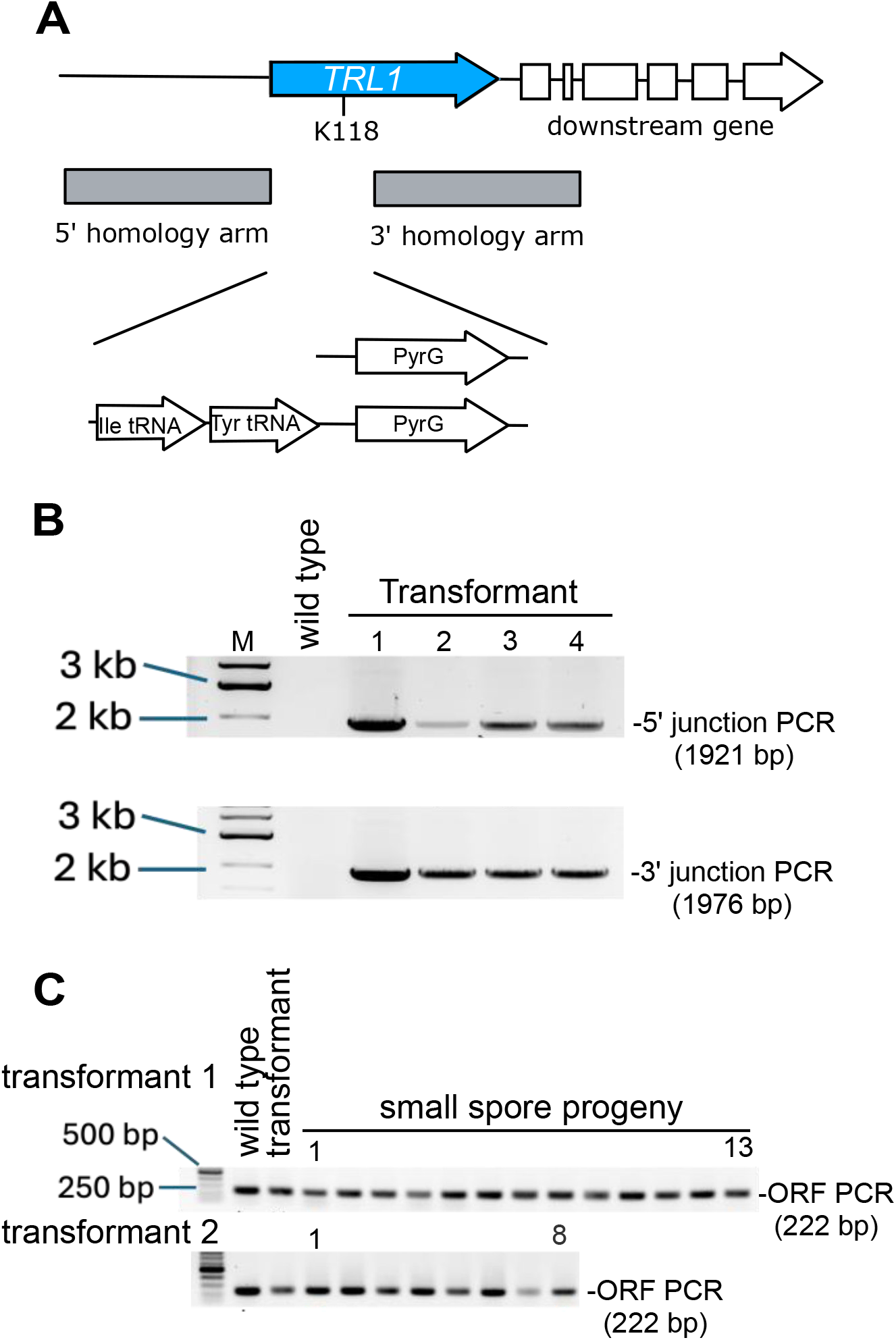
The ligase-only Trl1 of *Mucor circinelloides* is essential. (A) Schematic representation of the *trl1* locus of *M. circinelloides*. PCR primers flanking used in panels B to D are indicated with arrow heads. Black: primers flanking trl1, blue: primers internal to trl1. Homology arms to replace the start of the coding sequence, including the start codon and the active site lysine are indicated. The end of the coding sequence was left intact to not disturb expression of the downstream gene. The flanking primers are outside these homology arms. (B) *trl11* was disrupted by *pyrG.* Correct integration was confirmed by PCR with primers upstream of the homology and within *pyrG* (orange arrowheads) and with primers downstream of the homology and within *pyrG* (lower panel). (C) Small spores enriched for uni- and bi-nucleated spores were isolated and screened by PCR with primers within the coding region, showing that they all retained an intact *trl1* ORF. (D) *trl1* was disrupted by *pyrG* together with intronless versions of the Ile-UAU and Tyr-GUA tRNAs. Top: 3 transformants were confirmed by PCR as in B. Bottom: all small spores enriched for uni- and bi-nucleated spores retained an intact *trl1* ORF.

The *M. circinelloides* Trl1 is proposed to act in the splicing of two tRNAs (Tyr-GUA and Ile-UAU) and the *hac1* mRNA in the unfolded protein response (33, 35). The splicing of tRNAs should be essential because Tyr codons and AUA Ile codons cannot be translated in the absence of splicing. We therefore asked whether *M. circinelloides trl1* could be replaced by a *PyrG* cassette that also contains intron-less versions of the Tyr-GUA and Ile-UAU tRNAs. Again, we were able to obtain multiple heterokaryotic transformants, but small spore enrichment of mononucleated cells yielded only 36 strains that each contained an intact *trl1* gene (29 progeny from transformant 1, 3 progeny from transformant 2, and 4 progeny from transformant 3; Representative PCR results are shown in Figure 7D). These results indicate that expression of the two mature tRNAs fails to bypass *trl1Δ* in *M. circinelloides*. We observed that the *trl1lΔ::pyrG+2tRNAs* that express the intronless tRNAs grew slowly compared to the *trl1Δ::pyrG* heterokaryons, suggesting that removal of two tRNA introns has a detrimental effect, possibly because tRNA introns are required for tRNA modification (66, 67). Nonetheless, our failure to obtain homokaryotic *trl1Δ* small spores for five different transformants indicates that the atypical ligase of Mucorales is essential and may serve as a broad-spectrum antifungal target.

### All three catalytic activities of *Trypanosoma brucei* Trl1 are essential

Most parasites that infect humans also contain intron-containing tRNA genes and tRNA splicing enzymes. The tRNA ligase is either a RTCB ortholog (e.g. *Plasmodium* and *Giardia*) or a *TRL1* ortholog (e.g. *Trypanosoma* and *Leishmania*). To date, functional studies have only been carried out in *Trypanosoma brucei*, where inducible RNAi of *TRL1* is lethal (40). Importantly, this parasite has only one tRNA intron (in Tyr tRNA) and the growth defect due to inducible RNAi can be complemented by an intronless Tyr tRNA gene (40, 47). The *T. brucei* Trl1 also contains three distinct domains with conserved catalytic residues (Figure 8A). Thus, drugs that inhibit Trl1 might also be efficacious against trypanosomal infections if the inhibited domain is essential.

**Figure 8:**
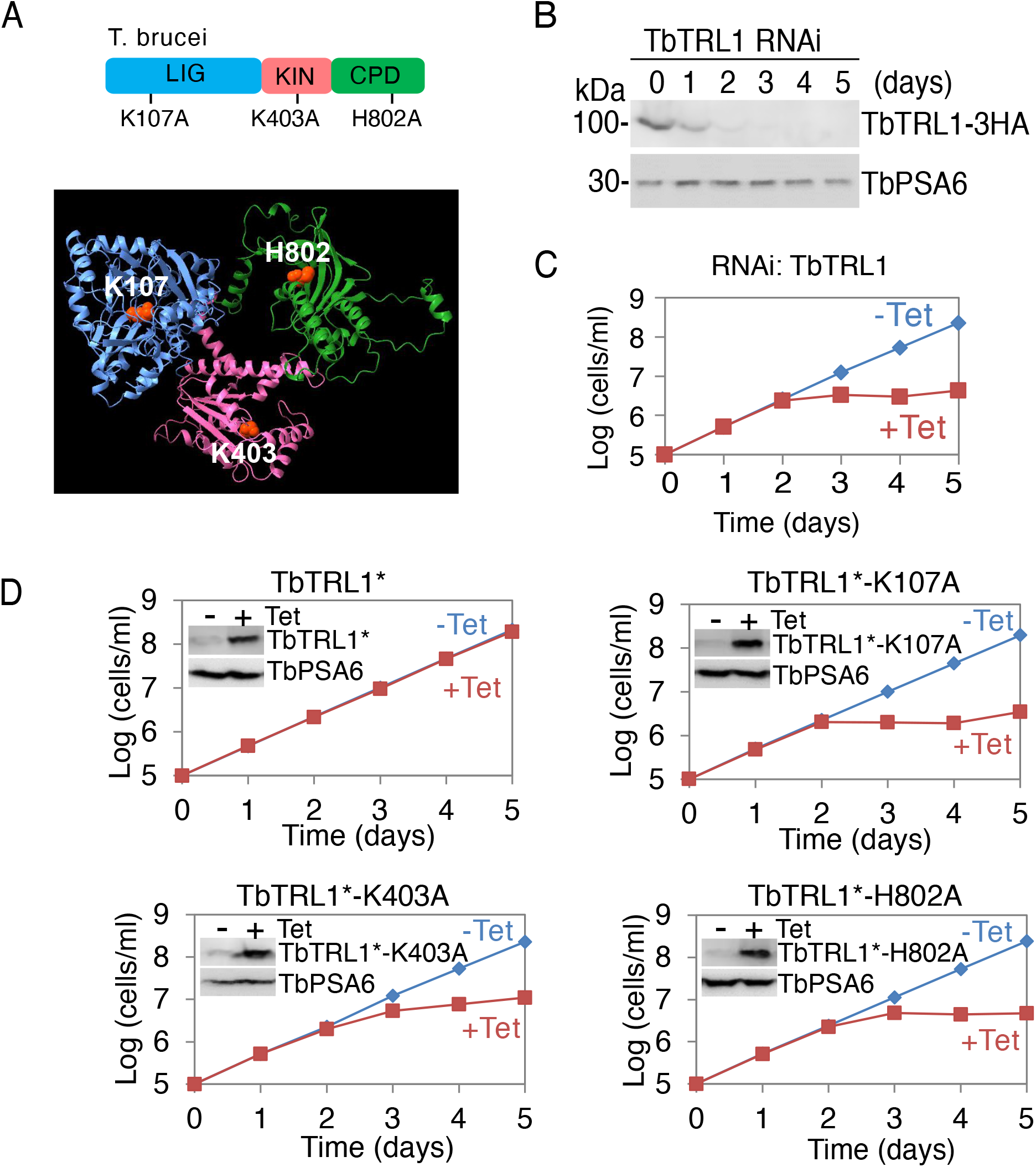
All three catalytic activities of *Trypanosoma brucei* Trl1 are essential. Schematic and AlphaFold predicted structure of *T. brucei* Trl1 with the three active sites indicated as in figure 4A. (B) Western blot analysis confirming depletion of *T. brucei* Trl1 following RNAi induction at different time points. Trl1 tagged with 3×HA (∼100 kDa) was detected using an anti-HA antibody. TbPSA6 served as a loading control. (C) RNAi-mediated depletion of *T. brucei* Trl1 following tetracycline induction caused a severe growth defect, whereas the non-induced control strain grew normally, confirming that *TRL1* is essential for the growth and viability of *T. brucei*. (D) The *T. brucei* Trl1 gene was partially recoded (*TRL1\**) to be resistant to the RNAi construct and the active sites were mutated. Expression of the wild-type Trl1 from the recoded construct complemented the RNAi phenotype, but expression of the recoded catalytically inactive *T. brucei* Trl1s did not.

We first confirmed that *TRL1* is essential using the same inducible RNAi approach reported before. Indeed, upon exposure to tetracycline, knockdown of *T. brucei TRL1* (Figure 8B) resulted in a pronounced growth defect beginning on day 3 after induction (Figure 8C). To determine which domains of *T. brucei TRL1* are essential, we generated ectopic overexpression constructs for wild-type *T. brucei TRL1* as well as mutants in each of the three domains. Expression of wild-type *TRL1* fully restored the growth defect observed upon inducible RNAi, indicating complete rescue of the silencing phenotype (Figure 8D). In contrast, cells expressing ligase (K107A), kinase (K403A), or CPD (H802A) domain catalytic mutants exhibited severe growth defects comparable to those observed in the uncomplemented RNAi line. These results indicate that all three domains of Trl1 are essential in *T. brucei* and represent potential drug targets for anti-trypanosomal drug development.

## DISCUSSION

The previous rationale for targeting Trl1 in antifungal drug development lies in the mechanistic differences from human RTCB, and the presence of Trl1 orthologs in pathogenic fungi (31, 38, 47, 68, 69). Based on this existing rationale, all three domains of Trl1 appeared to be valid antifungal drug targets; however, in vivo functional analysis of Trl1 has been largely limited to *S. cerevisiae*. From the *S. cerevisiae* studies, it was thought that fungal Trl1 enzymes contain three essential domains. Our results suggest that this may only be true for *S. cerevisiae* and very close relatives. Overall, our results suggest that the sealing domain of fungal Trl1 is essential, but that fungi generally contain redundant healing activities (Figure 9). Such a redundancy could be resolved in two ways that are exemplified by *Mucor* and *S. cerevisiae*. *Mucor* has lost the healing domains of Trl1, and presumably the alternative healing activities are now solely responsible for tRNA splicing (as well as *HAC1* splicing during the unfolded protein response) (Figure 9). *S. cerevisiae* appears to have resolved the redundancy instead by having lost the alternative healing activities and solely depending on the healing domains of Trl1. Unlike pathogenic fungi, the requirement of all three activities of Trl1 in *T. brucei* resembles that of *S. cerevisiae* Trl1. Therefore, each of the domains of *T. brucei* Trl1 appear to be viable drug targets, and it may be possible to develop a sealing domain inhibitor that has both anti-fungal and anti-trypanosome activities. This would negate some of the economic obstacles to the development of novel anti-trypanosomal drugs.

**Figure 9:**
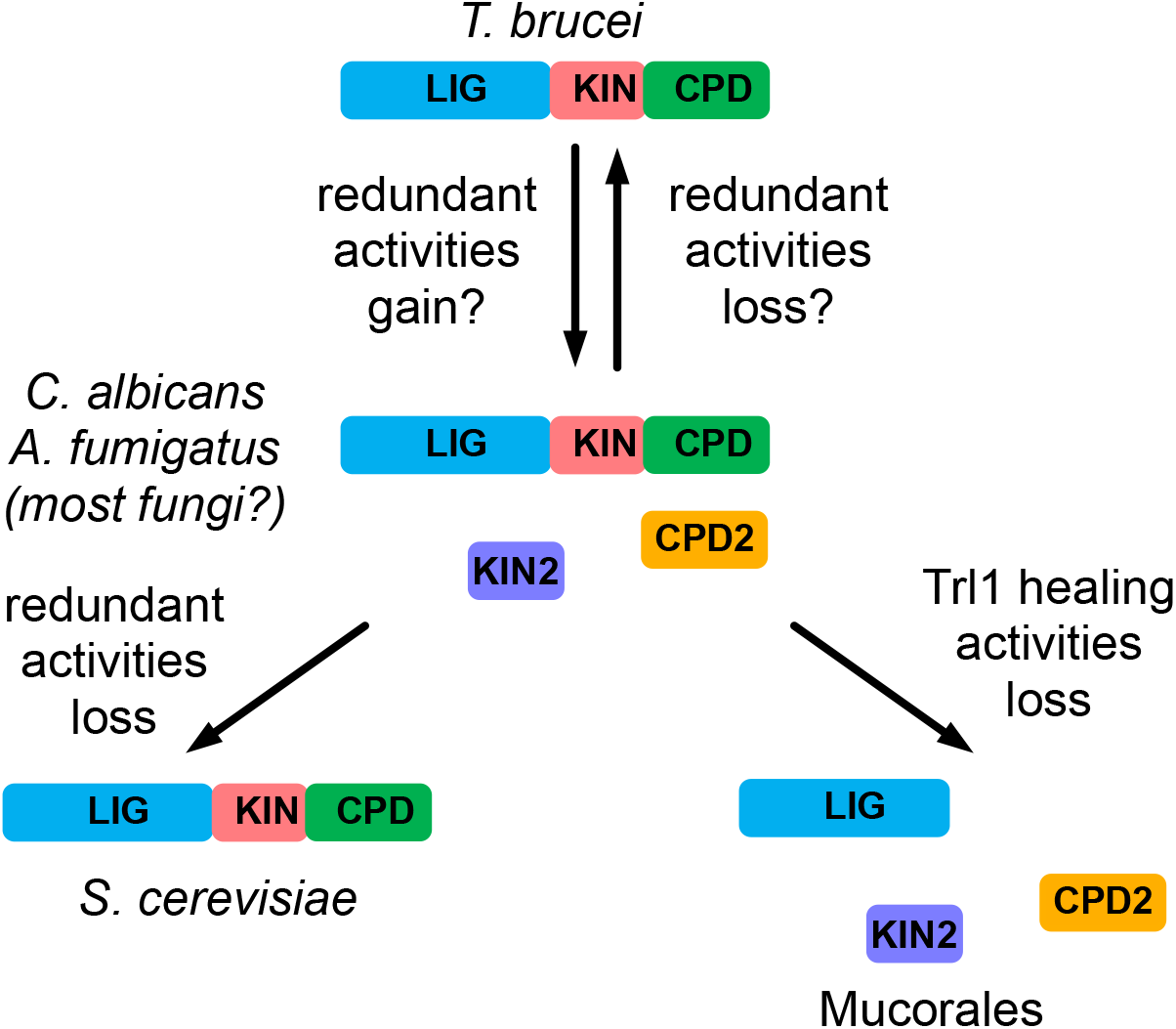
Model of TRL1 changes between organisms. Trl1 is depicted as in figure 3A. The unknown redundant enzymes are indicated as KIN2 and CPD2.

Importantly, our results indicate that the sealing domain of Trl1 is essential in pathogenic fungi. We have directly shown this for *C. albicans*, *A. fumigatus*, and *M. circinelloides*. Furthermore, at least in *S. cerevisiae* Trl1 inhibition leads to loss of viability, suggesting that a drug that inhibits the enzyme is likely to be fungicidal instead of merely fungistatic. We also show that *C. albicans* Trl1 sealing activity is required during an animal infection. A previous high-throughput screen suggested that *C. albicans* Trl1 is essential during a mouse infection (70), but the particular activity of Trl1 that was required was not identified. All of these results validate the sealing domain of Trl1 as a promising target for broad-spectrum antifungal drug development.

The sealing domain of Trl1 is distantly related to DNA ligases and RNA capping enzymes. The sealing domain itself consists of a catalytic core shared with these other enzymes and a Trl1-specific C-terminal subdomain. Despite the structural similarity in the catalytic core, the sequence similarity is low, and we expect that the divergence is sufficient to allow inhibitors to be specific. The catalytic core of Trl1 specifically recognizes the 2’ OH of RNA substrates (through R175 and H182). The Trl1-specific C-terminal extension is absent from DNA ligases and RNA capping enzymes but is important for RNA binding (46). It also confers a strict requirement for the 2’ monophosphate that is generated by the CPD domain (46). Notably, the CPD of Trl1 can be replaced in vivo by another CPD that generates a 2’ phosphate, but not the CPD of T4 PNK that generates a 3’ phosphate end, consistent with the strict requirement of a 2’ phosphate for the sealing step. These specificities of Trl1 for RNA and a 2’ phosphate, together with the very low sequence conservation, suggest there are drugable pockets that are absent from DNA ligases and RNA capping enzymes, and thus it should be possible to generate Trl1 specificity.

Although the Trl1 kinase mutants of *C. albicans* and *A. fumigatus* were viable, they consistently exhibit slower growth than the CPD mutant or wild-type Trl1-expressing cells. The growth defect of the kinase mutant of *A. fumigatus* Trl1 was very pronounced. The underlying cause of this growth phenotype remains unclear. One possibility is that the redundant kinase activity is suboptimal for tRNA splicing (and/or *HAC1* mRNA splicing). Alternatively, *S. cerevisiae* and plant Trl1 has been implicated in the kinase-mediated mRNA decay pathway, wherein phosphorylation of 5’ OH mRNA mediates their degradation by the 5’ monophosphate requiring RNase Xrn1 (71-73). Further studies are therefore required to elucidate why kinase mutants in pathogenic fungal species exhibit growth defects.

Our results indicate that fungi express alternative healing activities, but we have not identified them. Several enzymes with polynucleotide kinase domains are broadly conserved in fungi. This includes Clp1. Although yeast Clp1 is catalytically inactive, the human enzyme is active and can complement a Trl1-kinase defect when expressed in yeast. The inactivity of yeast Clp1 has been ascribed to the absence of a conserved aspartate and two conserved arginines (74). These inactivating mutations are shared across ascomycetes and basidiomycetes, but the Mucorales appear to encode an active Clp1 (with D121 R285 and K290 in the *Mucor circinelloides* protein HMPREF1544_09834). Therefore, *Mucor* Clp1 may be active in tRNA splicing. Mucor also encodes a second kinase that can participate in tRNA splicing in vitro and when expressed in *S. cerevisiae* (39). However, this protein is more closely related to DNA repair enzymes, including *S. pombe* Pnk1 (75-77), and its in vivo function remains to be determined. Finally, fungal genomes encode an ortholog of Grc3 (Nol9 in human), which is a polynucleotide kinase with a well-characterized function in rRNA 5’ end maturation (78, 79). Whether any of these polynucleotide kinases function in tRNA splicing in pathogenic fungi will require genetic analysis in these fungi. Alternatively, the redundant kinase activity might be provided by an unknown enzyme, and our results suggest it could be identified by looking for mutations synthetic lethal with Trl1 kinase deficiency. In contrast to the list of candidate polynucleotide kinases, there is a rarity of enzymes with CPD activity that produce 2’ monophosphate. Besides Trl1, only the rat CNP protein has been shown to have the capacity to complement a yeast defect in the CPD domain. The CPD domain belongs to the 2H family of phosphodiesterases. The 2H family has diverse activities and we were unable to identify fungal family members that are likely to act in tRNA splicing. Thus, while our work suggests that most fungi encode redundant tRNA exon healing activities, there are no close paralogs of Trl1, and the identity of those enzymes remains unknown. Regardless of the identity of the redundant healing activities, our results indicate that the sealing domain of Trl1 is a better drug target than the healing domains.

## MATERIALS AND METHODS

Plasmids and strains used are listed in Supplementary Table 1 and Supplementary Table 2.

### Structural analysis of fungal Trl1 orthologs

Structures of *C. albicans*, *A. fumigatus*, and *T. brucei* Trl1 orthologs were predicted using AlfaFold3 (80) and visualized using UCSF ChimeraX (81). Structures were aligned using the Matchmaker function of ChimeraX and catalytic residues were identified using structural and sequence alignment.

### *S. cerevisiae* plasmids, strains, and growth assays

Plasmids for the expression of *S. cerevisiae* Trl1 and the Trl1 orthologs from *C. albicans, A. fumigatus, and M. circinelloides* or *E. coli* RtcB in *S. cerevisiae* have been described previously (32). The RtcB coding region was codon optimized and CUG codons were removed. The plasmids containing point mutations in catalytic residues were generated either by amplifying the *TRL1* gene with mutagenesis primers and Gibson-assembled into pRS413 (32, 82), or using the QuikChange Lightning Site-Directed Mutagenesis Kit (Agilent Technologies). All plasmid constructs were verified by whole-plasmid nanopore sequencing (Plasmidsaurus).

The *S. cerevisiae trl1Δ* strain carrying a wild-type *TRL1* plasmid marked with the counter-selectable *URA3* marker has been described previously (32).

To express the wild-type or mutant *TRL1* genes, corresponding plasmids were transformed into the *trl1Δ* [*TRL1*, *URA3*] strain. Transformants were selected on synthetic complete (SC) medium lacking leucine and uracil (SC-Leu-Ura) or histidine and uracil (SC-His-Ura) (Sunrise Science). Plasmid-shuffle assays were performed by serial dilution and spot plating on medium containing 5-FOA, which selects for cells that have lost the wild-type *TRL1*/*URA3* plasmid (83). Growth of the *S. cerevisiae* strains was recorded after incubation for 4 days at 30°C.

The temperature-sensitive *S. cerevisiae trl1-ts* strain used to determine the fungicidal effect of Trl1 inhibition was previously described (54). It was grown in YPD at room temperature and then incubated at 37°C. Aliquots were taken during the 37°C incubation and plated onto YPD plates that were then incubated at room temperature.

### *C. albicans s*trains and growth assays

Conditional *TRL1* mutants of *C. albicans* were generated using the THE1 strain, a derivative of SC5314 that expresses the doxycycline-responsive transactivator (57). One *TRL1* allele was deleted using the SAT-FLP1 cassette from pSFS2 (56). Transformants were selected on YPD medium containing nourseothricin and correct integration was confirmed by PCR (yielding yAv4899).

To create a conditional expression allele, the promoter of the remaining *TRL1* allele in yAv4899 was replaced with the doxycycline-repressible TET-Off promoter (*tetO*) together with a *URA3* selectable marker. The promoter and *URA3* cassette was amplified from p97-CAU (57) and introduced by CRISPR-Cas9-mediated homologous recombination (58). Transformants were selected on SC-Ura and verified by PCR. Two independent creations of the conditional mutant had the same growth phenotype. The SAT1 marker from one resulting transformants was excised by growing on maltose-containing medium, and nourseothricin-sensitive colonies were isolated, creating yAV4901.

For complementation, the *C. albicans* wild-type or mutant *TRL1* coding sequences together with the native promoter and 3’UTR were cloned into the pSFBi-NEUT5L integration vector (a pSFS2 derivative) by Gibson assembly using the SacI and NotI restriction sites. Plasmids were linearized with StuI and electroporated into strain yAv4901 for targeted integration at the *NEUT5L* locus. Transformants were selected on YPD medium containing nourseothricin. Independent complemented isolates exhibited identical phenotypes. Multiple independently created isolates exhibited identical growth phenotypes. The genotypes of strain yAv4901 and the strains with wild-type or mutant *TRL1* or vector in the *NEU5L* locus were confirmed by whole-genome Nanopore sequencing performed at SeqCenter (Pittsburgh, PA). When indicated, the expression of the *TRL1* gene was repressed by growing cells in YPD medium supplemented with 100 µg/mL doxycycline.

For heterologous expression of *Escherichia coli* RtcB, the *rtcB* coding sequence was cloned into the CIp10-SAT1 vector (84) and integrated into the *RPS1* locus of yAV4901. Transformants were selected on YPD medium containing nourseothricin and verified by PCR.

Growth curves were generated by growing strains in biological triplicates in 96-well plates at 30°C in a Biotek Synergy H1 plate reader (Agilent) and measuring OD600 every hour. Plotted are the mean OD and the standard deviation of the three biological replicates.

To identify the two transposon insertions reporter in *TRL1*, the raw sequence data (28) was downloaded (SRA PRJNA490565), reads with the transposon end (GTATTTTACCGACCGTTACCGACCGTTTTCATCCCTA) were identified and trimmed with Cutadapt and mapped with Bowtie2 to the SC5413 reference genome.

### *A. fumigatus* strains and growth assays

To swap the endogenous *trl1* promoter with an *A. niger*-derived TET-on(PpkiA) promoter, a plasmid was created by assembling the 5’ flanking sequence of *A. fumigatus trl1,* the pyrithiamine resistance cassette, the TET-on(PpkiA) promoter sequence (85), and the 5’ coding sequences of *A. fumigatus trl1* in a pUC18 backbone using Gibson assembly (New England Biolabs). The resulting construct was transformed into *A. fumigatus* CEA17Δ*akub^KU80^* strain (86), and mutants were selected on AMM media supplemented with pyrithiamine.

To express the wild-type *A. fumigatus trl1+* in the TET-on(PpkiA) *A. fumigatus* conditional mutant, a plasmid was constructed containing the endogenous promoter/terminator of the *trl1* gene, genomic flanks for a safe haven region between AFUB_024480 and AFUB_024490, and a hygromycin resistance cassette (64). Linearized plasmid was transformed into the conditional mutant and selected on AMM media supplemented with pyrithiamine and hygromycin.

To express Trl1 with point mutations in the kinase and CPD domains, site-directed mutagenesis was performed using the Q5 site-directed mutagenesis kit (New England Biolabs). For the mutation in the sealing domain, a fragment containing the mutated site was synthesized by Eurofins Genomics and ligated into the linearized plasmid containing genomic flanks for a safe-haven region and a hygromycin resistance cassette. The resulting plasmids were linearized and transformed into the TET-on (PpkiA) *A. fumigatus* conditional mutant, and the transformants were selected on AMM media supplemented with pyrithiamine and hygromycin.

Correct genomic insertion of the TET-on promoter and *trl1* genes was confirmed by Southern blot (87).

For phenotyping assays, serial dilutions of spores were plated on AMM (88) and AMM supplemented with 3 µg/mL doxycycline, and strain growth was recorded after 48 hours of incubation at 37°C.

### TRL1 gene disruption in M circinelloides

For *M. circinelloides trl1* gene disruption, a plasmid was constructed by assembling 5’ and 3’ homology arms for the *trl1* gene flanking the *pyrG-dpl237* marker cassette amplified from the pSL13 plasmid (89). The space between the stop codon of *trl1* and the start codon of the downstream gene is only 97 bp. Thus, the 3’ homology arm was designed to delete the first 166 codons of the *trl1* ortholog (including the active site lysine) leaving 719 bp upstream of the downstream gene’s AUG intact. To replace the *M. circinelloides trl1* with intron-less Tyr and Ile UAU tRNA genes, a fragment for both intronless genes was synthesized (GenScript) and inserted into the above plasmid adjacent to the *pyrG-dpl237* marker cassette.

Gene deletion was performed as previously published (65, 90). Plasmids pAv2068 and pAv2069, carrying the *trl1Δ::pyrG or trl1Δ::pyrG+tRNAs* deletion cassettes, respectively, were linearized with SmaI, purified, and transformed into the MU402 (pyrG⁻) strain. Successful *trl1* disruption was confirmed by PCR with primers upstream of *trl1* (5’-TCATCCATCAACTGCCTCGA-3’), in the *pyrG* cassette (5’-ACCCACTCACTTTCCATTCG-3’ and 5’-TGCTTTTGTTGGCTGAGATG-3’) and downstream of *trl1* (5’-ACTGCGTTGTGTCATGATGG-3’). Progeny of small spores were screened by PCR with primers internal to *trl1* (5’-CGAGGGCTTTTCACCAAACA -3’ and 5’-TGGGTCTGTCTTCTCTGCAG-3’)

### RNAi and functional analysis in *T. brucei*

For *T. brucei* TRL1 gene knockdown, a Stem-Loop RNAi construct was generated by cloning a 600-bp DNA fragment of TbTRL1 gene (from position 720 to 1320) into the pSL-PAC plasmid (a derivative of pLew100). The resulting plasmid was used to transfect *T. brucei* strain 29-13, and transfectants were selected with 1.0 µg/ml puromycin. Cells were cloned by limiting dilution in a 96-well plate. To examine RNAi efficiency, TbTRL1 was endogenously tagged with a triple HA epitope in the TbTRL1 RNAi cell line and detected by Western blotting with the anti-HA antibody. RNAi was induced by incubating the RNAi cell line with 1.0 µg/ml tetracycline, and cell growth was monitored daily by counting cells with a hemacytometer.

To generate the TbTRL1 RNAi complementation cell line, the full-length TbTRL1, with the recoded sequence from position 1 to 1609 for RNAi resistance, was cloned into pLew100-3HA vector. The resulting plasmid, pLew100-TbTRL1*-3HA, was used to generate the K107A, K403A, and H802A mutants by site-directed mutagenesis. These plasmids were then each linearized by restriction digestion with NotI and used to transfect the TbTRL1 RNAi cell line. Transfectants were further selected with 1.0 µg/ml phleomycin and cloned by limiting dilution in 96-well plates. RNAi of TbTRL1 and ectopic overexpression of TbTRL1* and its three mutants were induced by incubating cells with 1.0 µg/ml tetracycline. Ectopic expression of wild-type and mutant TbTRL1*-3HA was detected by Western blotting with the anti-HA antibody. Cell growth was monitored daily by counting cells with a hemacytometer.

### *C. elegans* killing assay with Trl1 mutants

The methodology used to infect *C. elegans* with *C. albicans* in a liquid infection assay format was as previously described (91, 92), with a few modifications. Briefly, synchronized L4s (larval stage 4 *C. elegans*) were infected with the indicated *C. albicans* strains for 4 hours on BHI agar medium containing gentamycin (10 µg/ml) at 25°C. The nematodes were collected and washed four times with 2mL of sterile M9. They were collected by centrifugation at 750 x g for 30 seconds between each wash. The nematodes were then pipetted (∼30 per well, with two wells per condition, for a total of ∼60 worms assayed) into six-well plates containing 2 mL of liquid medium (20% BHI broth and 80% M9). Doxycycline was added to a concentration of 100 µg/ml at the indicated times. Plates were incubated at 25°C, and worm death was scored daily. Using GraphPad Prism (version 11.0), Kaplan-Meier survival curves were generated, and the log-rank test was used to compare them. P-values of <0.05 were considered to be statistically significant. At least three biological replicates were performed for all assays.

## ACKNOWLEDGEMENTS

This work was in part funded by a Dr. John J. Kopchick Research Award from The University of Texas MD Anderson Cancer Center UTHealth Graduate School of Biomedical Sciences to KSA, Ohio Eminent Scholar funds to AvH, NIH/NIAID Grant R01AI183606 to MCL and DAG, NIH/NIAID Grants R01AI143304 and R21AI147631 to MCL, a T32AI055449 fellowship to HBW, Federal Ministry for Research, Technology and Space (BMFTR: https://www.bmbf.de/), Germany, Project FKZ 01K12012 “RFIN—RNA-Biologie von Pilzinfektionen” to MGB, Project FKZ 01KI2515 “RFIN2.0–RNA-Biologie von Pilzinfektionen” to MGB, NIH/NIAID grants R01AI101437 and R01AI118736 to ZL, and NIH/NIAID grant R01AI182221 to SCL.

The funders had no role in the design of the study, data collection and analysis, the decision to publish, or the preparation of the manuscript.

We are grateful for John J. and Charlene Kopchick for their support of GSBS, Nathalie Seiler and Catherine Stuart for expert technical assistance and members of the van Hoof lab for valuable discussion.

