## Supplementa figure and tables for "Characterization of tRNA ligase function in pathogenic fungi and trypanosomes reveals the ligase domain as a promising drug target"

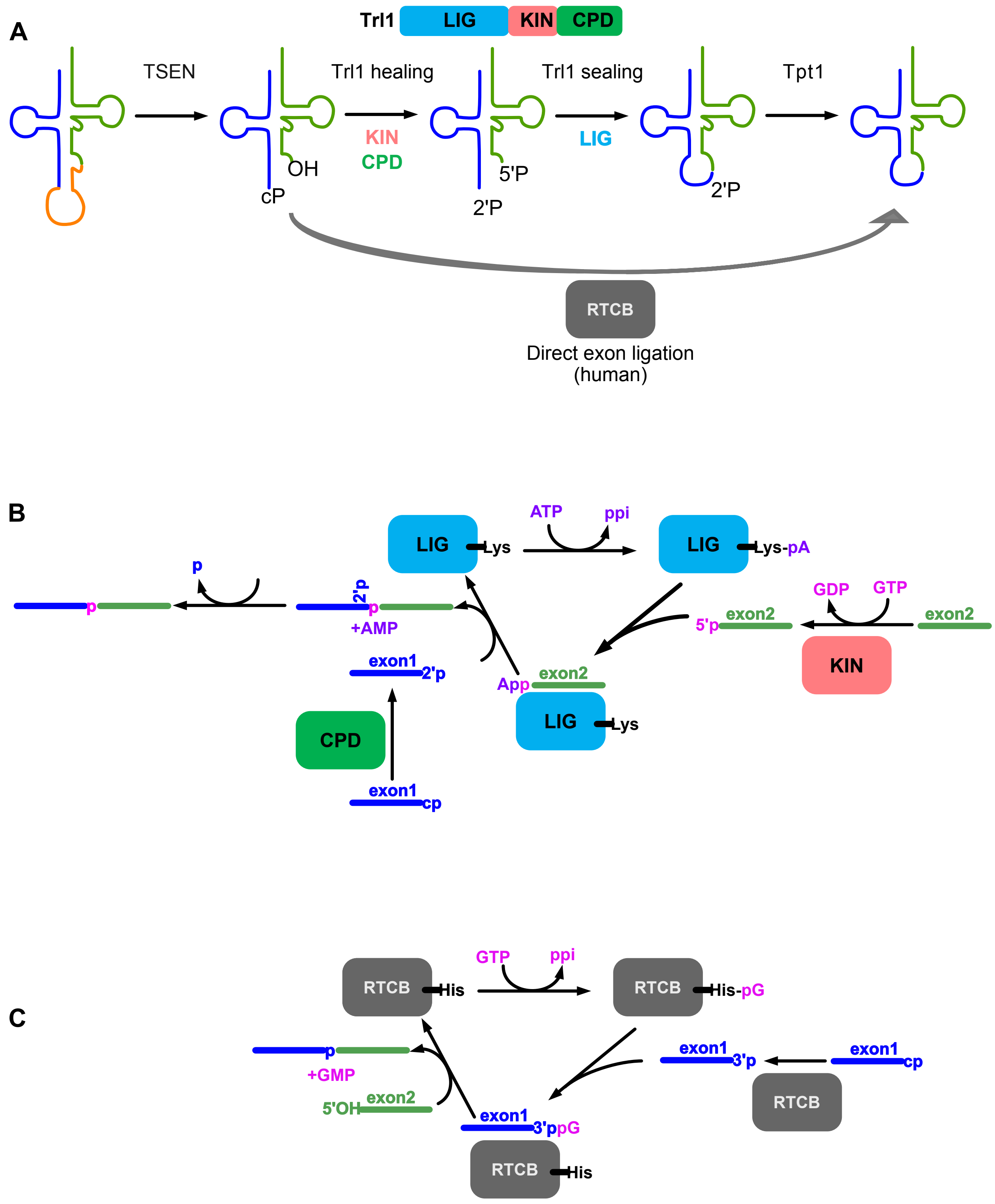


Supplemental figure

Supplemental figure: the tRNA splicing pathway in fungi and animals. (A) The overall pathway. (B) Detail of the fungal Trl1 pathway. (C) Detail of the fungal Trl1 pathway. The 5' exon is indicated in blue, the 3' exon in green, and the intron in orange. GTP is in pink, as is the phosphate in the phospodiester backbone of the spliced fungal tRNA, which is derived from the gamma phosphate of GTP. The phosphate indicated in blue is derived from the pre-tRNA and removed by Tpt1. This phosphate is released as ADP-ribose-1",2"-cyclic phosphate, which is omitted for simplicity.

Supplementary Table 1: Plasmids used in this study:

| Plasmid | Plasmid description | source |
|---|---|---|
| *Saccharomyces cerevisiae* plasmids | | |
| p425GPD | Empty vector, GPD promoter, *LEU2* marker | (1) |
| pRS413 | Empty vector, *HIS3* marker | (2) |
| pAv1511 | *Saccharomyces cerevisiae TRL1* in pRS416 | (3) |
| pAv1512 | *Saccharomyces cerevisiae TRL1* ORF p425GPD | (3) |
| pAv1655 | *Candida albicans TRL1* ORF in p425GPD | (3) |
| pAv1880 | *Aspergillus fumigatus TRL1* in p425GPD | (3) |
| pAv1514 | *Escherichia coli rtcB* ORF in p425GPD | This study |
| pAv1630 | *Saccharomyces cerevisiae* trl1-K114A in pRS413 | This study |
| pAv1614 | *Saccharomyces cerevisiae* trl1-K404A T405A in pRS413 | This study |
| pAv1616 | *Saccharomyces cerevisiae* trl1-H777A in pRS413 | This study |
| pAv1711 | *Saccharomyces cerevisiae* trl1-K404A T405A H777A in pRS413 | This study |
| pAv1970 | *Candida albicans trl1*-K108A in p425GPD | This study |
| pAv1971 | *Candida albicans trl1*-K425A T426A in p425GPD | This study |
| pAv1972 | *Candida albicans trl1*-H767A in p425GPD | This study |
| pAv2293 | *Aspergillus fumigatus trl1*-K117A in p425GPD | This study |

| | | |
|---|---|---|
| pAv2294 | *Aspergillus fumigatus trl1*-K397A T398A in p425GPD | This study |
| pAv2295 | *Aspergillus fumigatus trl1*-H775A in p425GPD | This study |
| Candida plasmids | | |
| pSFBI | *NEUT5L* integrative plasmid | |
| clp10-SAT1 | *RPS1* integrative plasmid | (4) |
| pAv2163 | *Candida albicans TRL1* in pSFBI | This study |
| pAv2163 | *Candida albicans trl1*-K108A in pSFBI | This study |
| pAv2163 | *Candida albicans trl1*-K425A T426A in pSFBI | This study |
| pAv2163 | *Candida albicans trl1*-H767A in pSFBI | This study |
| pAv2034 | *Candida albicans TRL1* in clp10-SAT1 | This study |
| pAv2075 | *Escherichia coli rtcB* coding sequence with *ACT1* promoter in clp10-SAT1 | This study |
| Aspergillus plasmids | | |
| pLS37 | *pkiA* and *ptrA* resistance with 5’ and 3’ homology flanks for AFUB_009140 (*trl1*) in pUC18 | This study |
| pLS33 | AFUB_009140 with endogenous promoter + terminator and *hph* resistance and 5’ and 3’ homology flanks for integration between AFUB_024480 and AFUB_024490 in pUC18 | This study |
| pLS35 | Same as pLS33 but point-mutations in AFUB_009140 (K397A T398A) | This study |
| pLS36 | Same as pLS33 but point-mutations in AFUB_009140 (H775A) | This study |

| | | |
|---|---|---|
| pLS39 | Same as pLS33 but point mutations in AFUB_009140 (K117A) | This study |
| Mucor circinelloides plasmids | | |
| pSL13 | PyrG-dpl237 | |
| pAv2068 | 5' and 3' homology arms of *Mucor circinelloides trl1* flanking *pyrG-dpl237* in pUC18 | this study |
| pAv2069 | 5' and 3' homology arms of *Mucor circinelloides trl1* flanking intronless Tyr (GUA) and Ile (UAU) tRNA and *pyrG-dpl237* in pUC18 | this study |
| Trypanosome plasmids | | |
| pSL-TbTRL1 | Stem-loop RNAi construct against TbTRL1 | this study |
| pLew100-TbTRL1* | Overexpression construct for expressing recoded TbTRL1 (TbTRL1*) | this study |
| pLew100-TbTRL1*-K107A | Overexpression construct for expressing recoded TbTRL1*-K107A mutant | this study |
| pLew100-TbTRL1*-K403A | Overexpression construct for expressing recoded TbTRL1*-K404A mutant | this study |
| pLew100-TbTRL1*-H802A | Overexpression construct for expressing recoded TbTRL1*-H802A mutant | this study |

| Supplementary Table 2: Strains used in this study: | |
|---|---|
| Strain | Genotype |
| *Saccharomyces* strains | |
| BY4741 | *matA, ura3-Δ0, leu2-Δ0, his3-Δ1, lys2-Δ0* |
| yAv4576 | *matA, ura3-Δ0, leu2-Δ0, his3-Δ1, lys2-Δ0, trl1Δ::NEO, [TRL1, URA3]* |
| yAv4360 | *matA, ura3-Δ0, leu2-Δ0, his3-Δ1, lys2-Δ0, trl1-ts* |
| *Candida* strains | |
| *THE1* | *ade2::hisG/ade2::hisG, ura3::imm434/ura3::imm434, ENO1/eno1::ENO1-tetR-ScHAP4AD-3xHA-ADE2* |
| yAv4899 | *ade2::hisG/ade2::hisG, ura3::imm434/ura3::imm434, ENO1/eno1::ENO1-tetR-ScHAP4AD-3xHA-ADE2, TRL1/trl1Δ* |
| yAv4901 | *ade2::hisG/ade2::hisG, ura3::imm434/ura3::imm434, ENO1/eno1::ENO1-tetR-ScHAP4AD-3xHA-ADE2, TRL1-URA3-TEToff/trl1Δ* |
| yAv5361 | *ade2::hisG/ade2::hisG, ura3::imm434/ura3::imm434, ENO1/eno1::ENO1-tetR-ScHAP4AD-3xHA-ADE2, TRL1-URA3-TEToff/trl1Δ, NEUT5L::SAT/NEUT5L* |
| yAv5355 | *ade2::hisG/ade2::hisG, ura3::imm434/ura3::imm434, ENO1/eno1::ENO1-tetR-ScHAP4AD-3xHA-ADE2, TRL1-URA3-TEToff/trl1Δ, NEUT5L::SAT-CalbTRL UTRs/NEUT5L* |
| yAv5349 | *ade2::hisG/ade2::hisG, ura3::imm434/ura3::imm434, ENO1/eno1::ENO1-tetR-ScHAP4AD-3xHA-ADE2, TRL1-URA3-TEToff/trl1Δ, NEUT5L::SAT-Calbtrl1-K108A/NEUT5L* |
| yAv5343 | *ade2::hisG/ade2::hisG, ura3::imm434/ura3::imm434, ENO1/eno1::ENO1-tetR-ScHAP4AD-3xHA-ADE2, TRL1-URA3-TEToff/trl1Δ, NEUT5L::SAT-Calb trl1-K425AT426A/NEUT5L* |

| | |
|---|---|
| yAv5337 | *ade2::hisG/ade2::hisG, ura3::imm434/ura3::imm434, ENO1/eno1::ENO1-tetR-ScHAP4AD-3xHA-ADE2, TRL1-URA3-TEToff/trl1Δ, NEUT5L::SAT-Calb trl1-H767A/NEUT5L* |
| yAv4903 | *ade2::hisG/ade2::hisG, ura3::imm434/ura3::imm434, ENO1/eno1::ENO1-tetR-ScHAP4AD-3xHA-ADE2, TRL1-URA3-TEToff/trl1Δ, rps1::vector/RPS1* |
| yAv4905 | *ade2::hisG/ade2::hisG, ura3::imm434/ura3::imm434, ENO1/eno1::ENO1-tetR-ScHAP4AD-3xHA-ADE2, TRL1-URA3-TEToff/trl1Δ, rps1::CalbTRL1*/RPS1 |
| | *ade2::hisG/ade2::hisG, ura3::imm434/ura3::imm434, ENO1/eno1::ENO1-tetR-ScHAP4AD-3xHA-ADE2, TRL1-URA3-TEToff/trl1Δ, rps1::rtcB/RPS1* |
| Aspergillus strains | |
| Wild type | *ΔakubKU80* |
| *trl1*$^{Tet\text{-}ON\text{(PpkiA)}}$ | *ΔakubKU80, trl1::ptrA::TET-on(PpkiA)* |
| *trl1*$^{Tet\text{-}ON\text{(PpkiA)}}$ *+trl1* | *ΔakubKU80, trl1::ptrA::TET-on(PpkiA), AFUB_024480::hph::trl1+* |
| *trl1*$^{Tet\text{-}ON\text{(PpkiA)}}$ *+trl1*$^{K397AT398A}$ | *ΔakubKU80, trl1::ptrA::TET-on(PpkiA), AFUB_024480::hph::trl1*$^{K397AT398A}$ |
| *trl1*$^{Tet\text{-}ON\text{(PpkiA)}}$ *+trl1*$^{H775A}$ | *ΔakubKU80, trl1::ptrA::TET-on(PpkiA), AFUB_024480::hph::trl1*$^{H775A}$ |
| *trl1*$^{Tet\text{-}ON\text{(PpkiA)}}$ *+trl1*$^{K117A}$ | *ΔakubKU80, trl1::ptrA::TET-on(PpkiA), AFUB_024480::hph::trl1*$^{K117A}$ |
| Mucor strains | |
| CBS277.49 | |

1. D. Mumberg, R. Muller, M. Funk, Yeast vectors for the controlled expression of heterologous proteins in different genetic backgrounds. *Gene* **156**, 119–122 (1995).
2. R. S. Sikorski, P. Hieter, A system of shuttle vectors and yeast host strains designed for efficient manipulation of DNA in Saccharomyces cerevisiae. *Genetics* **122**, 19–27 (1989).
3. K. S. Ahammed, A. van Hoof, Fungi of the order Mucorales express a "sealing-only" tRNA ligase. *RNA* **30**, 354–366 (2024).
4. A. M. Murad, P. R. Lee, I. D. Broadbent, C. J. Barelle, A. J. Brown, CIp10, an efficient and convenient integrating vector for Candida albicans. *Yeast* **16**, 325–327 (2000).
